# A streamlined proof of concept CRISPR-based environmental biosurveillance platform for the detection of closely related invasive rats

**DOI:** 10.64898/2026.09.24.753988

**Authors:** Benjamín Durán-Vinet, Antoinette J. Piaggio, Nicholas Foster, Anna Clark, Madison Salyer, Gert-Jan Jeunen, Jackson Treece, Sara Ferreira, Catherine Collins, Stacey Buckelew, Neil J. Gemmell

## Abstract

Environmental biosurveillance requires technologies that can rapidly translate genomic information into sensitive, specific, and deployable detection tools. Here, we present the first proof-of-concept application of SENTINEL (Smart Environmental Nucleic-acid Tracking using Inference from Neural-networks for Early-warning Localization), integrating artificial-intelligence-guided target discovery, isothermal amplification, CRISPR-based detection, and environmental DNA. Using Rattus exulans, R. norvegicus, and R. rattus as a challenging model of closely related invasive vertebrates, SENTINEL achieved species-specific discrimination without detectable cross-reactivity and sensitivities of 10 aM for plasmid DNA and 10 fg µL^⁻¹^ for genomic DNA. Selected assays also showed reproducible semi-quantitative performance and successful eDNA validation. This first application of CRISPR-based environmental biosurveillance to terrestrial invasive vertebrates demonstrates a transferable framework for converting sequence data into field-deployable assays, with potential applications across biosecurity, conservation, agriculture, and environmental monitoring.

**Graphical abstract:** 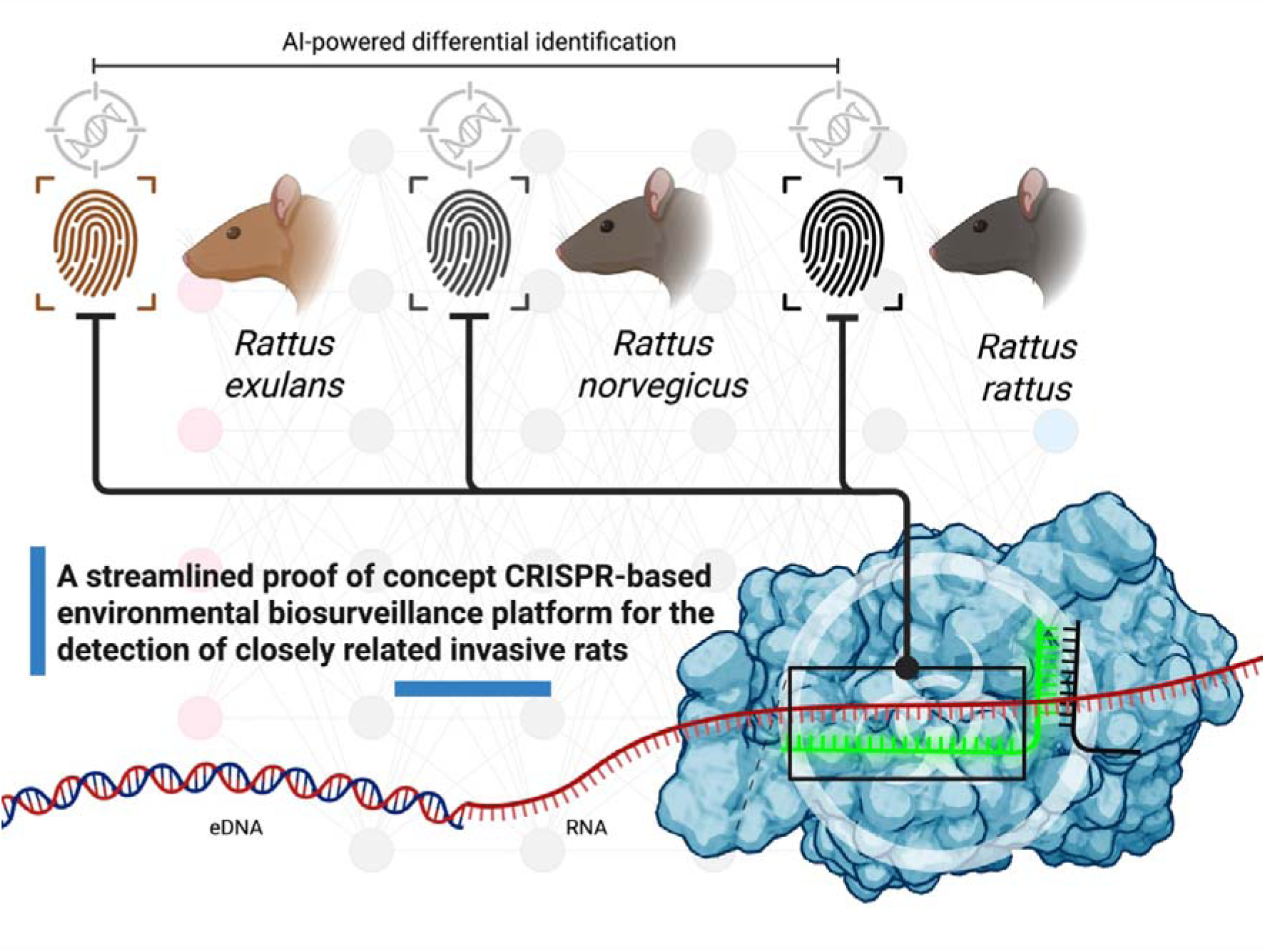

## Introduction

Invasive rat species are among the most widespread and damaging mammalian pests globally, exerting severe ecological, biosecurity, and economic impacts (Davis et al. 2023), costing the US alone US$27 billion each year (Richardson et al. 2025). *Rattus norvegicus* (brown rat) and *Rattus rattus* (black rat) have been unintentionally introduced to various ecosystems worldwide, with *R. norvegicus* and *R. rattus* achieving near-global distributions (Puckett et al. 2020; Richardson et al. 2025; Stokes et al. 2009). *Rattus exulans* (Pacific rat) was likely intentionally transported from South East Asia with people as they settled the islands of the Pacific (Roberts 1991), and today is distributed across the Pacific. Due to these differing histories, attitudes and values about *Rattus* differ, with *R. exulans* being culturally significant to some Pacific cultures (Wehi et al. 2021).

Rats’ omnivorous diets, rapid reproduction, and behavioural adaptability enable them to outcompete native species and decimate biodiversity (Graham et al. 2024; Honzák et al. 2024). On islands, rats are leading drivers of avian declines (Lee et al. 2022), and that can lead to complete collapses of native fauna (Moore et al. 2022), making early detection essential for effective biosecurity management, eradication efficiency, and long-term ecosystem recovery (Lieurance et al. 2023; Ringler et al. 2023). Accordingly, the eradication of invasive rats is increasingly being prioritised and implemented to conserve and recover biodiversity and mitigate economic impacts. An early, proactive commitment to invasive species management can be substantially different from a delayed management approach, with costs that could escalate to hundreds of billions if actions are delayed (Brondízio et al. 2019; Cuthbert et al. 2022).

Traditional monitoring methods, including camera surveys or rodent detection dogs (Davis et al. 2023), provide valuable information but can lack sensitivity, are labour-intensive, spatially limited, and slower (Steibl et al. 2025). Because biosecurity systems increasingly require rapid, scalable, portable, and highly sensitive tools, molecular approaches have emerged as powerful complementary strategies to traditional methods (Piaggio et al. 2025; Williams et al. 2025). These molecular approaches leverage the detection of environmental nucleic acids (eNAs/eDNA); eNAs provide a transformative solution by enabling the detection of species from trace genetic material shed into the environment. The use of eDNA can capture presence signals from hair, saliva, urine, skin cells, or faeces without requiring direct observation or physical capture of individuals (Bass et al. 2023; Kelly et al. 2024; Steibl et al. 2025). This capability is particularly advantageous for cryptic or elusive species such as invasive rodents (Piaggio et al. 2025), where early detection can greatly influence the success of eradication programmes (Lieurance et al. 2023). Quantitative polymerase chain reaction (qPCR) is widely used for eDNA detection across various targets (Cangelosi et al. 2024; Guri et al. 2024; Piaggio et al. 2025), although it faces challenges related to inhibition from environmental matrices and the need for high portability in point-of-use (POU) deployments (Williams et al. 2021, 2023; Yang et al. 2024). Although novel portable qPCR systems are now commercially available (e.g.,(Kona et al. 2025)), isothermal amplification technologies require fewer components to be functional, which could lead to more portable, cost-effective, and lightweight devices (e.g., (Wang et al. 2024)).

To implement an isothermal assay, clustered regularly interspaced short palindromic repeats (CRISPR) and CRISPR-associated proteins (Cas) have emerged in the eNAs field, offering a next-generation solution to the POU portability and inhibition challenges. Cas effector proteins (e.g., Cas13) can be programmed with guide RNAs (gRNAs) to recognise specific sequences with high fidelity (Abudayeeh and Gootenberg 2021; Gootenberg et al. 2018, 2017; Kellner et al. 2019). Upon target binding, these nucleases exert a *trans*-collateral cleavage activity over specific reporters on a probe that harbours a short sequence of nucleotides flanked by a fluorescent dye and a quencher. This has been harnessed and coupled with isothermal amplification technologies (e.g., recombinase polymerase amplification – RPA) to increase sensitivity, while Cas effectors provide an additional layer of specificity. Therefore, when this probe is degraded during sample incubation, a signal that can be detected rapidly (Abudayeeh and Gootenberg 2021; Gootenberg et al. 2018), even in the presence of low-abundant targets (de Puig et al. 2021; Yang et al. 2024). The application of this combination of CRISPR-Cas and eNAs for ecological, environmental biosurveillance and biosecurity applications has been termed CRISPR-based environmental biosurveillance (CRISPR-eBx; (Durán-Vinet et al. 2025a)). To date, no study has applied CRISPR-eBx to detect invasive terrestrial species and differentiate between closely related invasive target species.

Therefore, in this study, we have deployed our previously developed SENTINEL platform (Smart Environmental Nucleic-acid Tracking using Inference from Neural-networks for Early-warning Localization; (Durán-Vinet et al. 2025)), an artificial intelligence–powered, Cas13-based CRISPR-eBx platform, for the detection of three closely related invasive rats: *R. exulans*, *R. norvegicus* and *R. rattus*. SENTINEL integrates a pre-trained deep learning model (Ackerman et al. 2020; Metsky et al. 2022) for *in silico* guide-primer pair discovery (GPP; a set of primers flanking a gRNA), to predict and rank highly active combinations tailored to custom user input parameters, which can depend on assay objectives, including species-level resolution, tolerance to sequence biodiversity variability and coverage. Cas13 was selected due to its straightforward, one-pot application compatible with pre-amplification, whereas Cas12 *trans*-collateral activity is directed against single-stranded DNA (ssDNA), which can naturally degrade ssDNA primers (Chen et al. 2018). Workarounds do exist, but they add extra operational steps (Cheng et al. 2025).

Our study aimed to deploy the SENTINEL platform across *Rattus* spp. mitochondrial genomes and to experimentally validate *in silico* predicted GPPs for the differential identification of *R. exulans*, *R. norvegicus*, and *R. rattus* using genomic DNA (gDNA). To achieve this, we validated our targets on plasmid DNA and then we used gDNA reference samples from both the United States (US) and New Zealand (NZ) as a proof of concept for an assay capable of detecting these reference populations while exhibiting no off-target activity against closely related species.

After validation in gDNA, our study successfully validated eDNA results for *R. exulans* and *R. rattus*, demonstrating that the SENTINEL platform is an accessible, reproducible, and streamlined CRISPR-eBx platform, enabling swift and specific biosurveillance of highly similar invasive species. Our results further demonstrate that AI integration can strongly support and streamline environmental biosecurity of known and emerging threats.

## Material and methods

This study can be summarized in three major steps (**Figure**): (1) artificial intelligence deployment for guide-primer pair discovery, (2) SENTINEL functionality validation and (3) SENTINEL differential identification panel validation.

### Reference whole mitochondrial genome retrieval

Full mitochondrial genomes were used as the reference target discovery sequences for the SENTINEL assay development targeting *Rattus exulans* (Rexu, Pacific rat), *R. norvegicus* (Rnor, brown rat) and *R. rattus* (Rrat, black rat). As reference mitochondrial genomes, we retrieved *R. exulans* (GenBank accession number NC_012389), *R. norvegicus* (NC_001665) and *R. rattus* (NC_012374).

Each reference sequence was used in GenBank megaBLAST (version 2.14.0; (Camacho et al. 2009)) to retrieve further sequences with at least ≥90.0% pairwise identity to the target sequence of the respective invasive species. Sequences were retrieved following this criterion for *R. exulans (n = 32)*, *R. norvegicus (n = 65)* and *R. rattus (n = 45)*, respectively. Accession numbers for each retrieved are listed in **Supplemental Table 1**.

Retrieved sequences for each species were aligned using the Multiple Alignment using Fast Fourier Transform (MAFFT; (Nakamura et al. 2018)). High-coverage regions and polymorphic sites were identified with a 50% minimum sequence frequency threshold on the consensus sequence for genomic reference. This means that for either, a potential polymorphic site or a high-coverage region, the region was required to be present on ≥50% across the retrieved sequences. Then, we generated consensus sequences for all alignments with a 0% majority call threshold (most common bases are called in each site to build a consensus to avoid ambiguities in plasmid inserts. These steps are crucial for correctly assessing and understanding polymorphic sites and the high-coverage region landscape for target species. All *in silico* analyses, such as guide-primer pair exploration and curation, were performed in Geneious Prime (2024.0.5; https://www.geneious.com).

### Artificial intelligence deployment for guide-primer pair design

We used ADAPT (Activity-informed Design with All-inclusive Patrolling of Targets, version 1.6.0; https://adapt.run/) (Metsky et al. 2022), and previous work experience for the deployment of the SENTINEL platform (Ackerman et al. 2020; Durán-Vinet et al. 2025) on closely related invasive rat species. Briefly, ADAPT is a pre-trained, end-to-end artificial intelligence that designs highly active guide-primer pairs (GPPs; a spacer sequence and flanking RPA primers) from input FASTA sequences.

For each species (*R. exulans*, *R. norvegicus* and *R. rattus*), an alignment of mitochondrial genomes were used as "on-target sequences" in FASTA format while incorporating identified potential off-target sequences as "excluded sequences" in FASTA format (Supplemental Table 1). For each target species, the sequences were run and optimised independently of each other via ADAPT. Off-target sequences were identified as sequences with full complementarity – 100% pairwise identity using GenBank MegaBLAST using obtained spacer sequences after a first exploratory ADAPT run without the ***‘--specific-against-fastas***’ command. Identified off-target accession numbers were added as a single, unaligned FASTA file in a second final run (**Supplemental Table 1**). Additional parameters, including primer length, primer GC content, spacer length, spacer specificity, and other customizations, were applied to optimise LwaCas13a crRNA modelling and diagnostic performance prediction for SENTINEL as explained in the original ADAPT article (Metsky et al. 2022), and reference GitHub repositories (https://github.com/broadinstitute/adapt; https://github.com/bduranvinet/SENTINEL-ed).

The set of commands to accomplish these tasks are provided **(Supplemental Table 2).** The output of ADAPT was a .TSV file containing the predicted activity, the crRNA spacer sequence (28 nucleotides), and the RPA primer sequences (30 nucleotides each). Only the spacer portion of the crRNA was checked with megaBLAST, and full specificity was enforced (no mismatches allowed). RPA in the SENTINEL platform is used to enrich target DNA, increasing sensitivity, while LwaCas13a provides the specificity and detection layer. Therefore, RPA specificity is not heavily enforced in our approach, but rather heavily enforced on LwaCas13a spacer sequence.

Selected detection sites were used for gene synthesis and custom plasmid construction (GenScript, US, custom product). Plasmids harbouring these custom inserts were used as proof of concept for functionality prior to further assay development into gDNA. Sequences are available in Supplemental Table 3.

### Genomic DNA reference samples for cross-reactivity and sensitivity testing

The gDNA samples from US *Rattu*s species (target species and other closely related rodents to *Rattus* spp), *Neotoma albigula,* and *Mus musculus* used in this study were provided by the United States Department of Agriculture (USDA). The gDNA samples from NZ *Rattus* species and *Mus musculus* were provided by University of Otago collaborators (see **Supplemental Table 4**). All gDNA samples were aliquoted and stored at -20°C.

When more than one individual of the same species was available, gDNA samples of the same species were pooled in equal amounts into 0.1 ng uL^-1^ final concentration stocks. For gDNA sensitivity, 0.1 ng µL^-1^ (-1), 0.01 ng µL^-1^ (-2), 0.001 ng µL^-1^ (-3), 0.0001 ng µL^-1^ (-4) and 0.00001 ng µL^-1^ (-5) inputs were tested. The final concentrations in the SENTINEL platform assays (2µL sample input in final 20 µL reaction, 1:10 dilution of input) for each dilution were 10,000 fg uL^-1^ (-1), 1,000 fg uL^-1^ (-2), 100 fg uL^-1^ (-3), 10 fg uL^-1^ (-4) (see **Supplemental Table 5**). For all cross-reactivity assays, a fixed input value of 0.1 ng µL^-1^ was used. All gDNA extractions were quantified with a Qubit™ 4 Fluorometer (ThermoFisher Scientific, US) using Qubit™ 1X dsDNA High Sensitivity (ThermoFisher Scientific, US) following the manufacturer’s instructions.

To select a single and final GPP candidate for each species, each GPP was evaluated independently, with selection decisions made on a case-by-case basis according to its individual experimental performance. All GPPs in this study were screened using 100pM plasmid DNA input with the target insert complementary to the corresponding GPP. Moreover, selected best-performing GPP, gDNA functionality was also screened at 0.1 ng uL^-1^ gDNA input in parallel with rat reference samples from United States (US) and New Zealand (NZ) for all GPPs.

### SENTINEL fluorescence assays

The Cas13 mastermix with Cas13a optimized reaction buffer was produced as previously reported (Durán-Vinet et al. 2025). The mastermix composition for a single RPA pellet (yields four reactions) is as follows: 24.78 µL of RNase-free water (Invitrogen, US, #10977015), 16.80 µL of 5X optimized Cas13a reaction buffer, 2.10 µL of Murine RNAse inhibitor (New England Biolabs - NEB, US, #M0314S), 3.36 µL of ribonucleotide mix (NEB, US, #N0466S), 4.20 µL of 10 µM reporter (poly-U_5_, GenScript, US, custom), 4.20 µL of 9.6 µM pre-pooled forward and reverse primer (GenScript, US, custom), 3.36 µL of T7 RNA polymerase (NEB, US, #M0251S), 8.40 µL of 500 nM LwaCas13a (GenScript, US, #Z03486; diluted in RNAse-free water and 1X RNAse-free PBS pH 7.4, Invitrogen, US, #AM9624), 4.20 µL of 1 µM crRNA (GenScript, US, custom) and 4.20 µL of 280 mM MgOAc (TwistDx, US). All components were added in the described order. We accounted for a 5% pipetting error when scaling up the reactions. All mastermixes were freshly produced and used immediately.

A final volume of 75.60 µL of mastermix was slowly mixed into one RPA pellet (TwistDx, US). 18 µL of mastermix was then aliquoted in wells of a white 8-tube strip while using a 96-well adaptor (Roche Diagnostics, US, LightCycler® 8-Tube Strip Adapter Plate, #06612598001), and then 2 µL of sample was added for plasmid and gDNA testing. Then, 8-tube white strips were spun down for 15 seconds at full speed in a minifuge and immediately run in LC480 II LightCycler qPCR systems (Roche Diagnostics, US) at 39°C, for one hour with data acquisitions every 60 seconds in the FAM channel (excitation at λ465 nm and detection at λ510 nm) as previously reported (Durán-Vinet et al. 2025). RNAse-free water was used as a negative control in all assays unless otherwise specified. All assays were run in triplicate plus a negative non-target control per RPA pellet. All data is reported as a noise-corrected, baseline-subtracted fluorescence, unless otherwise specified.

### SENTINEL platform environmental DNA proof of concept

When deploying candidate GPPs on the SENTINEL platform, we used 4 µL of each eDNA sample. The previous protocol was therefore adjusted in the following components: 16.38 µL of RNase-free water (Invitrogen, US, #10977015). The final volume of the mastermix was 67.20 µL, and 16 µL of mastermix was aliquoted into wells of a white 8-tube strip using a 96-well adaptor. All other components and running parameters were maintained. Accordingly, we obtained samples from US (Wake Island) and NZ (place) that were benchmarked with qPCR assays for *R. exulans* and *R. rattus*, respectively. *R. norvegicus* eDNA samples were not available for validation at the time of this study. True negative eDNA samples used in this study came from a rat-free US island but with a population of invasive *Mus musculus* using qPCR benchmark (Oh et al. 2021; Piaggio et al. 2025; Midway Island, US). All environmental DNA samples information used in this study are provided in Supplemental Table 6.

### Statistics and SENTINEL metrics

Statistical analyses, including t-tests, ANOVA, post-hoc multiple comparisons, and correlation analyses, were performed using Prism 10 (Version 10.4.0 for MacOS, GraphPad Software, Boston, Massachusetts USA, www.graphpad.com). Differences between two GPPs were assessed using Welch’s t-test. For comparisons involving multiple groups, one-way ANOVA followed by Dunnett’s multiple-comparisons test was used to compare each dilution group mean endpoint fluorescence with the control to observe significant, i.e., positivity.

The t-test and *p*-values are given in parentheses when corresponding. All data plots (heatmap, column plots, and curves) were created with Prism 10. All illustrations were created with BioRender (www.biorender.com/). Fold change was calculated as endpoint fluorescence divided by the average of the last five timepoints of the endpoint baseline fluorescence, and the standard deviation was then added to the average. Relative fold change was only used for SENTINEL’s GPPs validation. For all statistical tests, a significance level of α = 0.05 was used. The obtained *p-*values in text are presented with three significant figures, while figures display extended *p*-values.

Values normalization was done using the highest values from gDNA for each independent set of GPPs for each species.

To further evaluate SENTINEL deployment and CRISPR-eBx performance more broadly, an important question remains: how can GPP performance be empirically compared beyond fluorescence intensity alone? To address this, we developed a quantitative metric termed activation time (A_t_), which represents the earliest time point at which the signal generated by a given GPP–target combination can be reliably distinguished from the negative control. This provides an additional measure of GPP performance based on assay kinetics, complementing comparisons based solely on fluorescence magnitude (see Supplemental methods for further information).

## Results

### ADAPT pipeline identified at least one guide-primer pair candidate in Rattus spp. for species-specific identification

The ADAPT pipeline provided one, seventeen, and three GPPs for *R. exulans*, *R. norvegicus,* and *R. rattus*, respectively. The two GPPs with the highest expected activity and on-target coverage, when possible, were selected for experimental screening (**Table 1**).

**Table 1.** Obtained candidate guide-primer pair for invasive rat species.

| Target species | Genomic target hit | Guide-primer pair ID | Length (bp) | Expected activity | On-target coverage | Number of on target sequences |
| --- | --- | --- | --- | --- | --- | --- |
| *Rattus exulans* | ND1 | Rexu-GPP-3188 | 121 | 3.19 | 1.00 | 32 |
| *Rattus norvegicus* | 16S | Rnor-GPP-1555 | 106 | 3.30 | 0.99 | 65 |
|  | ND5 | Rnor-GPP - 13385 | 89 | 3.55 | 0.99 |  |
| *Rattus rattus* | ND2 | Rrat-GPP-4487 | 114 | 3.02 | 1.00 | 45 |
|  | COX3+tRNA-Gly+ND3 | Rrat-GPP-9426 | 212 | 3.09 | 0.98 |  |

Abbreviations: GPP: guide-primer pair; Rexu: *Rattus exulans*; Rnor: *Rattus norvegicus*; Rrat: *Rattus rattus*.

### Validation of SENTINEL with candidate guide-primer pairs using plasmid DNA and genomic DNA

In order to select a single GPP candidate for each species, all GPPs were validated in an independent, case-by-case basis process. On *R. exulans*, Rexu-GPP-3188 showcased good functionality at 100 pM plasmid input with an average A_t_ = 4.57 min (**Figure 2A; Supplemental Table 7**). Moreover, Rexu-GPP-3188 showcased a significantly higher endpoint fluorescence on US *R. exulans* gDNA in comparison to NZ *R. exulans* (**Figure 2B**). Moreover, Rexu-GPP-3188 showcased A_t_ versus US and NZ gDNA equal to 16.39 and 19.54 min, respectively. Therefore, Rexu-GPP3188 was selected as candidate GPP for further sensitivity screening in plasmid DNA and gDNA. An arbitrary positivity threshold was set at a conservative F_thr_ ≍ 1.872 (equal to 3-fold change units; Supplemental Table 7).

**Figure 1.**
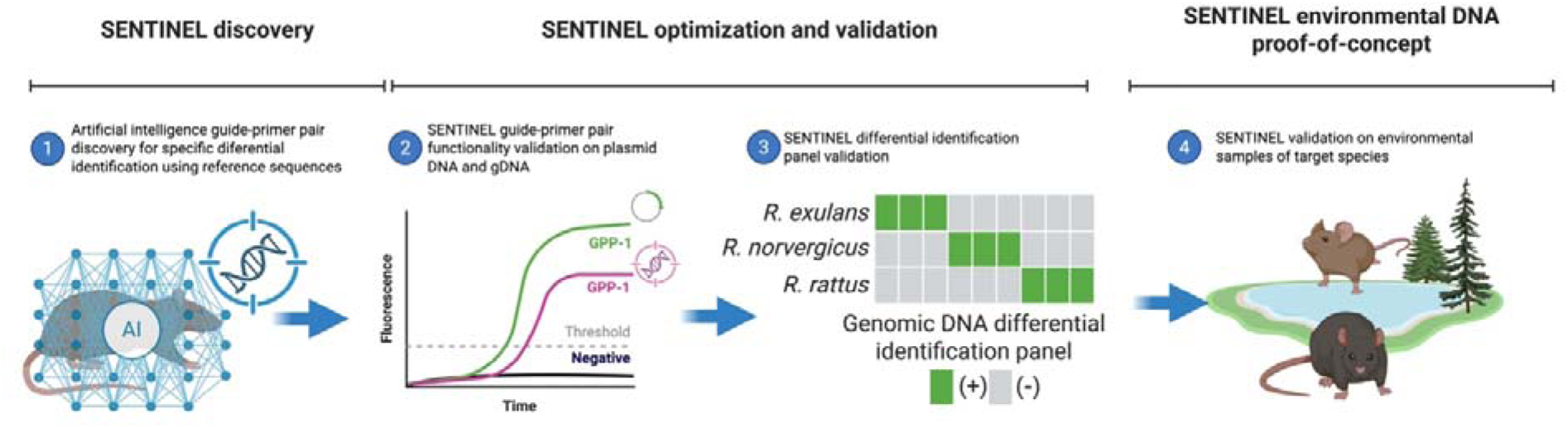
SENTINEL workflow for species-specific invasive rat differential identification. (1) Artificial intelligence guide-primer pair discovery. (2) SENTINEL validation on plasmid and gDNA. (3) SENTINEL differential identification panel validation using gDNA reference of target rat species. (4) SENTINEL validation as a proof of concept in environmental samples. gDNA: Genomic DNA; GPP: guide-primer pair; SENTINEL: Smart Environmental Nucleic-acid Tracking using Inference from Neural-networks for Early-warning Localization.

**Figure 2.**
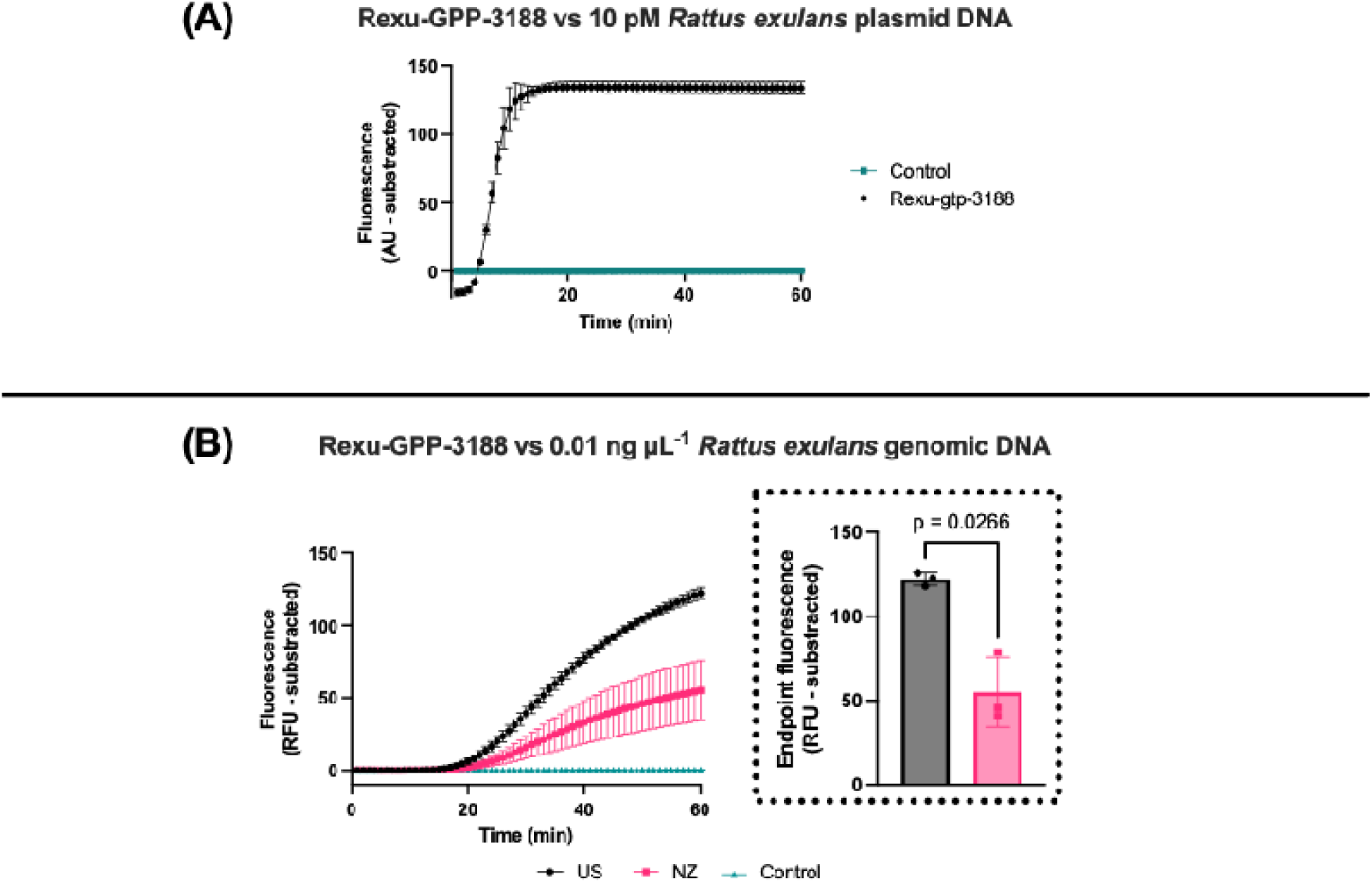
SENTINEL Rexu-GPP-3188 validation in plasmid and genomic DNA. **(A)** Rexu-GPP-3188 assay versus 10 pM *R. exulans* plasmid DNA. **(B)** Rexu-GPP-3188 assays versus 0.01 ng µL-1 R. exulans gDNA from United States (US) and New Zealand (NZ). Endpoint fluorescence is significantly higher in US gDNA (t-test; t = 5.523; *p* = 0.027). All data points were obtained with n = 3. Columns within the dotted black boxes represent the subtracted endpoint fluorescence mean; all error bars show ± SD (n = 3). Abbreviations: gDNA: genomic DNA; GPP: guide-primer pair; RFU: Relative fluorescence units; Rexu: *Rattus exulans*; SENTINEL: Smart Environmental Nucleic-acid Tracking using Inference from Neural-networks for Early-warning Localization.

For *R. norvegicus*, the results showed that both GPPs (Rnor-GPP-1555 and Rnor-GPP-13385) had no significant fluorescence when tested on 10 pM plasmid DNA (t-test, t = 1.675, *p* = 0.239; **Figure 3A**). Mean A_t_ from Rnor-GPP-1555 and Rnor-GPP-13385 was 3.20 and 1.65 min, respectively **(Supplemental Table 7)**. When further tested, there was a significant difference in endpoint fluorescence performance between Rnor-GPP-1555 vs Rnor-GPP-13385 on 0.01 ng µL^-1^ of gDNA from US reference *R. norvegicus* sample (t-test, t = 4.932 *p* = 0.022; **Figure 3B**). Additionally, the mean A_t_ for US *R. norvegicus* gDNA for Rnor-GPP-1555 and Rnor-GPP-13385 was equal to 16.55 and 12.51 min. Jointly from these results, Rnor-GPP-13385 was selected as the most active GPP candidate for *R. norvegicus* detection using a conservative F_thr_ ≍ 1.872 (equal to ∼3.0 fold-change units). Further screening was done with the selected candidate Rnor-GPP-13385 to assess activity over NZ *R. norvegicus* reference sample. Results indicated that Rnor-GPP-13385 had better endpoint fluorescence (t test, t = 5.148, *p* = 0.017, **Figure 3C**) and activation time for NZ *R. norvegicus* gDNA.

**Figure 3.**
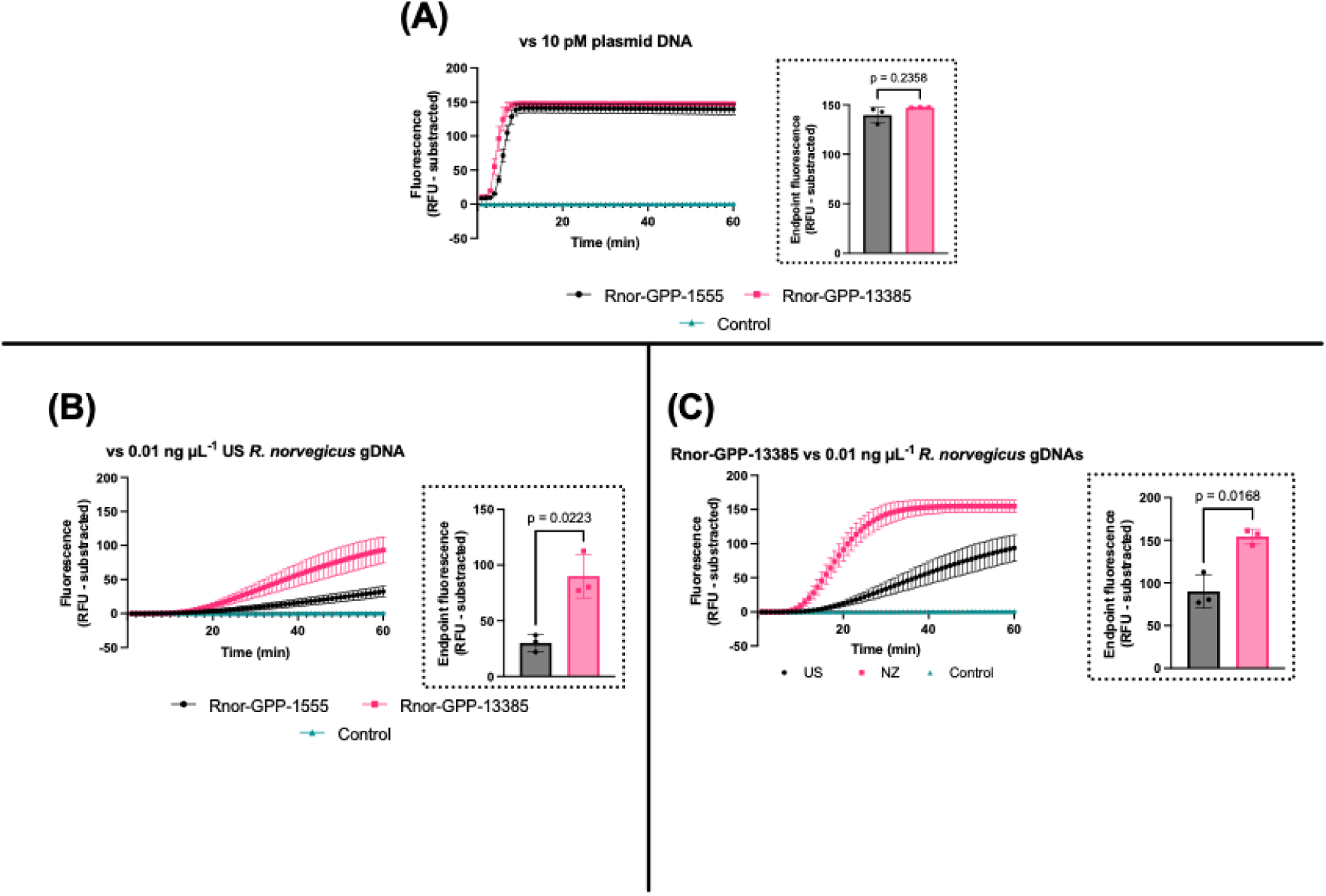
SENTINEL guide-primer pair screening on *Rattus norvegicus*. **(A)** SENTINEL activity on 10 pM of plasmid insert harbouring *R. norvegicus* target GPP sites. Black outlined dotted box shows Rnor-GPP-1555 and Rnor-GPP-13385 subtracted endpoint fluorescence with no significant different in endpoint fluorescence (t-test, t = 1.675 *p* = 0.239). **(B)** SENTINEL activity on 0.01 ng µL^-1^ US *R. norvegicus* gDNA. Black outlined dotted box shows Rnor-GPP-1555 and Rnor-GPP-13385 endpoint fluorescence in gDNA from US *R. norvegicus* reference sample with a significant difference (t-test, t = 4.932, *p* = 0.022). (C) SENTINEL Rnor-GPP-13385 activity over US and NZ R. norvegicus gDNA. Black outlined dotted box shows Rnor-GPP-13385 endpoint fluorescence in gDNA from US and NZ *R. norvegicus* gDNA reference samples with a significant difference (t-test, t = 5.148, *p* = 0.017). All data points were obtained with n = 3. Columns within the black boxes represent the subtracted endpoint fluorescence mean; error bars show ± SD (n = 3). Abbreviations: gDNA: genomic DNA; GPP: guide-primer pair; RFU: Relative fluorescence units; Rnor: *Rattus norvegicus*; SENTINEL: Smart Environmental Nucleic-acid Tracking using Inference from Neural-networks for Early-warning Localization.

For *R. rattus*, the results shown that Rrat-GPP-4487 had a significant difference versus Rrat-GPP-9426 in endpoint fluorescence (t-test, t = 32.72, *p* < 0.001; **Figure 4A**). Moreover, Rrat-GPP-4487 and Rrat-GPP-9426 had mean A_t_ equal to 4.16 and 2.49 min using a F_thr_ ≍ 1.508 (equal to ∼8.0 fold-change units), respectively (**Supplemental Table 7**). Following tests showcased that Rrat-GPP-9426 had a higher endpoint fluorescence on 0.01 ng µL^-1^ of gDNA from US reference *R. rattus* sample (t-test, t = 7.349, *p* = 0.017; **Figure 4B**) Additionally, the mean A_t_ for US *R. rattus* gDNA for Rrat-GPP-4487 and Rrat-GPP-9426 was equal to 13.91 and 7.14 min. Further screening was done with the selected candidate Rrat-GPP-9426 to assess activity over NZ *R. rattus* reference gDNA sample. We found that Rrat-GPP-9426 detection efficiency for NZ *R. rattus* was not significant versus US *R. rattus* gDNA (t-test, t = 1.654, *p* = 0.236; **Figure 4C**). However, Rrat-GPP-9426 showed faster A_t_ values for both US and NZ equal to 7.14 and 7.92 min (**Supplemental Table 7**). Hence, Rrat-GPP-9426 was selected as candidate GPP for *R. rattus* detection.

**Figure 4.**
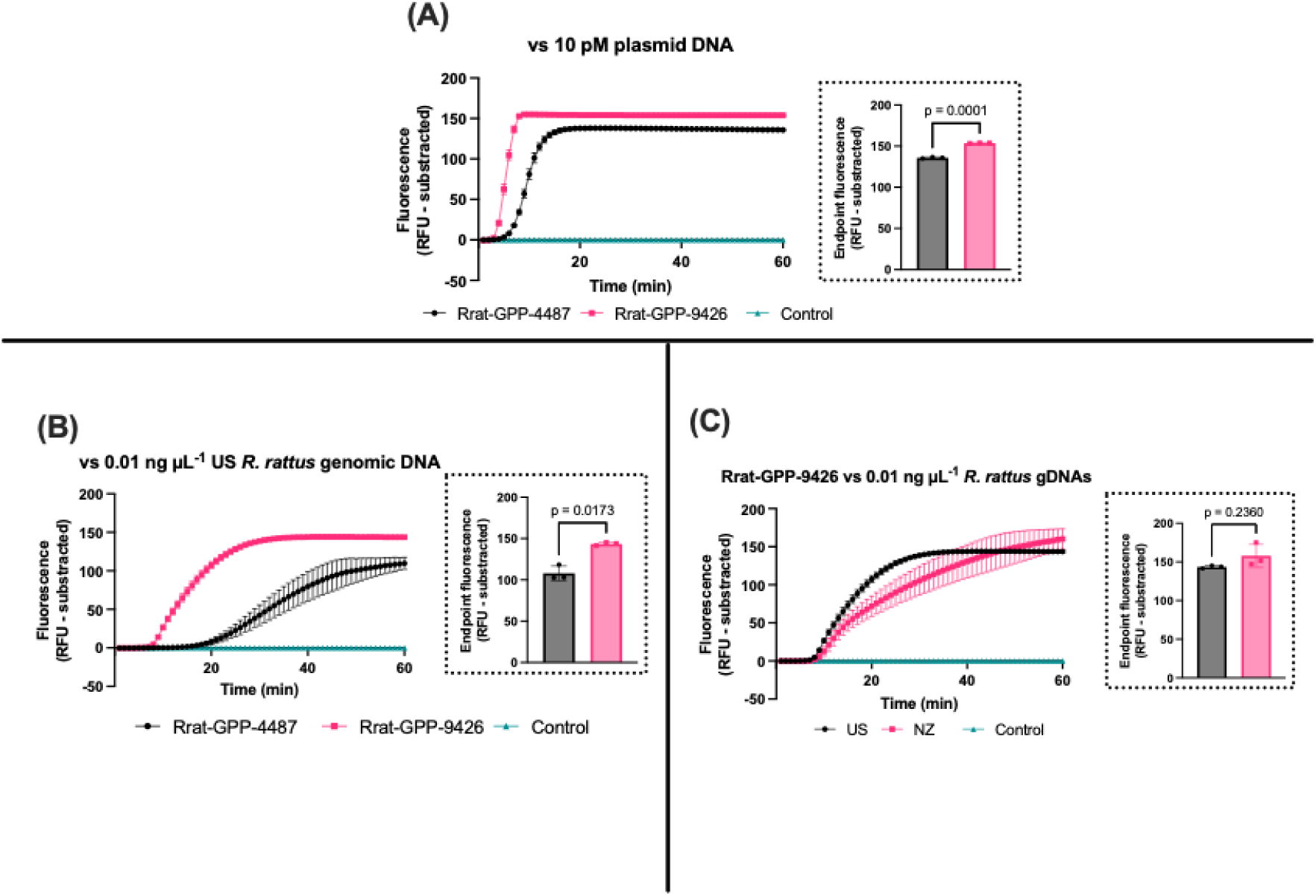
SENTINEL guide-primer pair screening on *Rattus rattus*. **(A**) SENTINEL activity on 10 pM of plasmid insert harbouring *R. rattus* target GPP sites. Black outlined dotted box shows Rrat-GPP-4487 and Rrat-GPP-9426 endpoint fluorescence with a significant difference (t = 32.72, *p* **(B)** SENTINEL GPP activity on 0.01 ng µL^-1^ US *R. rattus* gDNA reference sample. Black outlined dotted box shows Rrat-GPP-4487 and Rrat-GPP-9426 subtracted endpoint fluorescence test on gDNA from US *R. norvegicus* reference sample (t = 7.349, p = 0.013, p < 0.05). (C) SENTINEL Rrat-GPP-9426 activity over US and NZ *R. rattus* gDNA. Black outlined dotted box shows Rrat-GPP-9426 endpoint fluorescence in gDNA from US and NZ *R. rattus* gDNA reference samples with a significant difference (t-test, t = 1.654, *p* = 0.236). All data points were obtained with n = 3. Columns represent means; error bars show ± SD (n = 3). Abbreviations: gDNA: genomic DNA; GPP: guide-primer pair; Rrat: *Rattus rattus*. RFU: relative fluorescence units; SENTINEL: Smart Environmental Nucleic-acid Tracking using Inference from Neural-networks for Early-warning Localization.

To facilitate downstream sensitivity screening on gDNA, we only used the best-performing gDNA reference samples for sensitivity assays, i.e., highest endpoint fluorescence and fastest A_t_. In detail, for Rexu-GPP-3188 was further tested with US gDNA, Rnor-gpp-13385 was further tested with NZ gDNA and Rrat-GPP-9426 was further tested with US gDNA reference samples.

### SENTINEL candidate guide-primer pair cross-reactivity performance showcase highly specific differential identification of closely related invasive rat species

Candidates GPPs were tested against a panel composed of several closely related rats Error! Reference source not found. human and other rodents at 0.01 ng µL^-1^ gDNA (**Supplemental Table 4)**. Overall, no GPP had a significant fold change across all GPP across all seven species (**Figure 5**)

**Figure 5.**
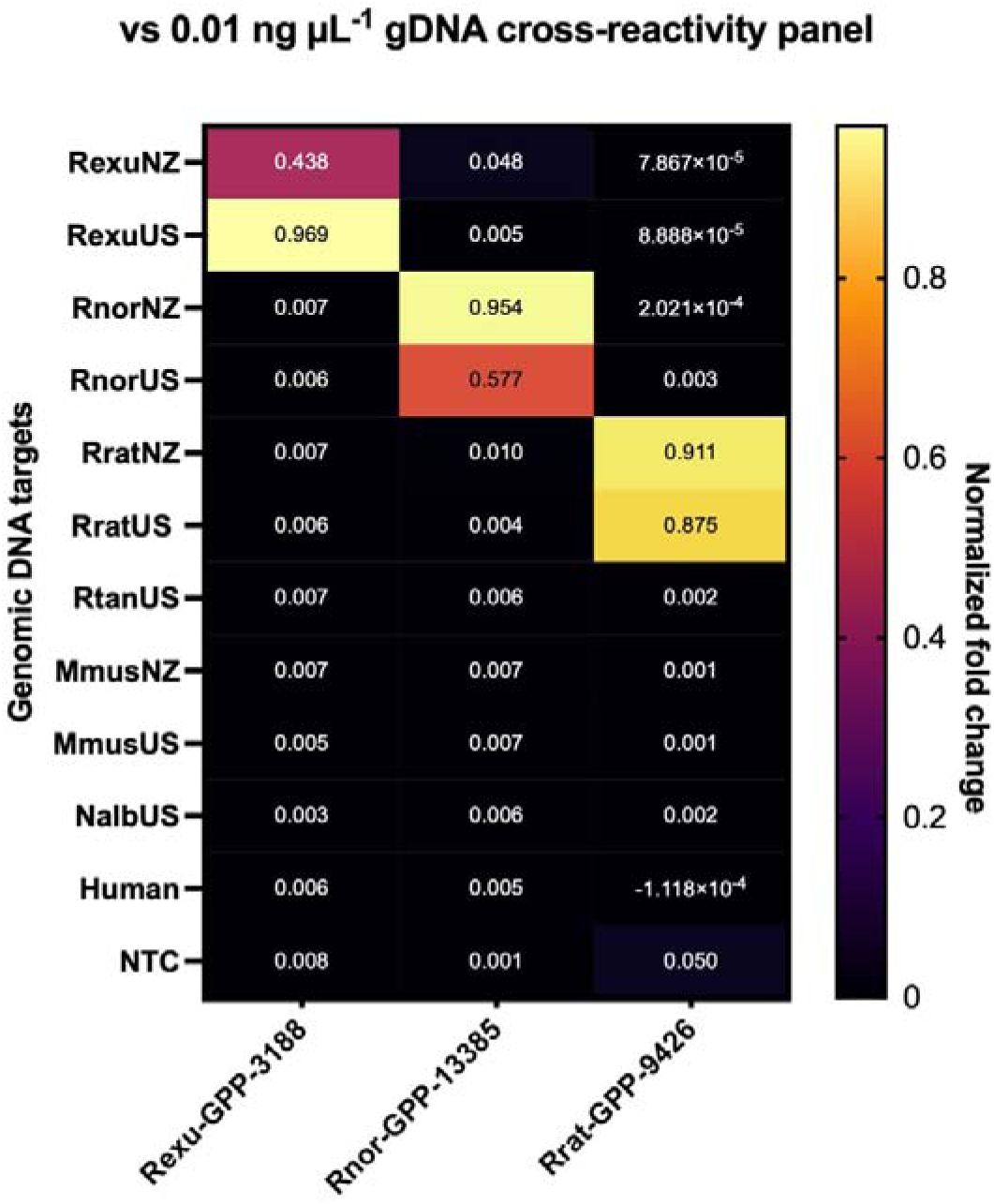
SENTINEL all candidate GPPs cross-reactivity panel. All species were tested in triplicate with a final concentration of 0.01 ng µL^-1^. All boxes represent the mean (n = 3). Abbreviations: GPP: guide-primer pair; Mmus: *Mus musculus*; Nalb: *Neotoma albigula*; Rexu: *Rattus exulans*; Rnor: *Rattus norvegicus*; Rrat: *Rattus rattus*; Rtan: *Rattus tanezumi*; NTC: non-template control; SENTINEL: Smart Environmental Nucleic-acid Tracking using Inference from Neural-networks for Early-warning Localization.

### SENTINEL candidates showcase semi-quantification capabilities in plasmid DNA and genomic DNA of invasive rat species

We explored whether the selected GPPs maintained previously reported semi-quantitative capabilities of SENTINEL (Durán-Vinet et al. 2025). All GPPs — Rexu-GPP-3188, Rnor-GPP-13385 and Rrat-GPP-9426 — showcased a significant fold change on plasmid DNA versus the negative controls (**Figure 6**). Rexu-GPP-3188 fold-change on plasmid DNA using a one-way ANOVA was significant (F(3, 8) = 597, *p* < 0.05). Homogeneity of variance was confirmed (Brown–Forsythe: F(3, 8) = 1.930, *p* = 0.203). Dunnett’s post-hoc test, comparing each dilution against the negative control, indicated that all detected dilutions were significantly different from negative control (adjusted p < 0.05; **Figure 6A**). Rnor-GPP13385 analyzed with a one-way ANOVA was also significant (F(3,8) = 4842, *p* <0.05). Homogeneity of variance was confirmed (Brown–Forsythe: F(3, 8) = 1.440 *p* = 0.302). Dunnett’s post-hoc test, comparing each dilution against the negative control, indicated that all dilutions were significantly different (adjusted *p* < 0.05; **Figure 6B**). Finally, Rrat-GPP-9426 results analyzed with a one-way ANOVA showed a significant effect (F(3, 8) = 943.2, *p* < 0.05). Homogeneity of variance was met (Brown–Forsythe: F(3, 8) = 1.679, *p* = 0.248) Dunnett’s post-hoc test, comparing each dilution against the negative control, indicated that all dilutions were significantly different (adjusted *p* < 0.05; **Figure 6C**). Semi-quantification capabilities were demonstrated with dilutions of plasmid DNA for *R. exulans*, *R. norvegicus* and *R. rattus* were R^2^ = 0.73, 0.76 and 0.72, respectively (**Figure 6**).

**Figure 6.**
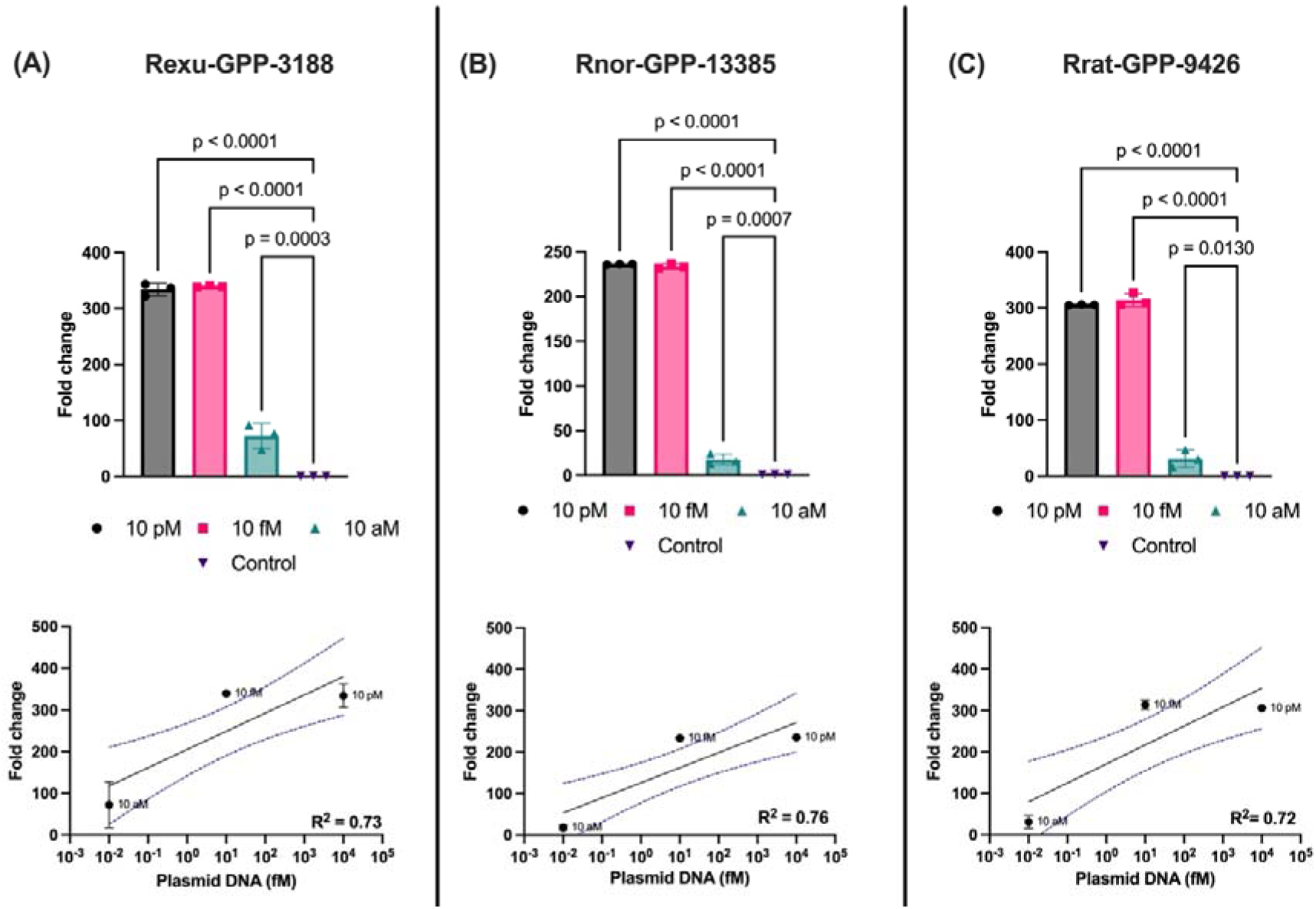
SENTINEL guide-primer pair semi-quantitative window in plasmid DNA. **(A)** Rexu-GPP-3188 fold change on plasmid DNA one-way ANOVA was significant, and Dunnett’s post-hoc test, comparing each dilution against the negative control, indicated that all GPPs were significantly different (adjusted *p* < 0.05). **(B)** Rnor-GPP-13385 fold change on plasmid DNA one-way ANOVA was significant and Dunnett’s post-hoc test, comparing each dilution against the negative control, indicated that all GPPs were significantly different (adjusted *p* < 0.05) **(C)** Rrat-GPP-9426 fold change on plasmid DNA one-way ANOVA was significant and Dunnett’s post-hoc test, comparing each dilution against the negative control, indicated that all GPPs were significantly different (adjusted *p* < 0.05). All dilutions were labelled accordingly in the interpolation graphs (10 aM, 10 fM and 10 pM). Abbreviations: GPP: guide-primer pair; Rexu: *Rattus exulans*; Rnor: *Rattus norvegicus*; Rrat: *Rattus rattus*; SENTINEL: Smart Environmental Nucleic-acid Tracking using Inference from Neural-networks for Early-warning Localization.

Then, to further challenge SENTINEL, we tested candidate GPPs with a more complex sample: gDNA. All GPPs showcased a significant fold change in gDNA versus the negative controls (**Figure 7**). Rexu-GPP-3188 one-way ANOVA showcased a significant fold change in gDNA (F(4, 10) = 79.56, *p* < 0.05). Homogeneity of variance was confirmed (Brown–Forsythe: F(4, 10) = 0.891, p = 0.503). Dunnett’s post-hoc test, comparing each dilution against the negative control, indicated that all detected dilutions were significantly different from the negative control (adjusted p < 0.05; **Figure 7A**). Rnor-GPP-13385 one-way ANOVA results were also significant (F(4,10) = 274.3, *p* <0.05). Homogeneity of variance was confirmed (Brown–Forsythe: F(4, 10) = 0.735 p = 0.589). Dunnett’s post-hoc test, comparing each dilution against the negative control, indicated that all dilutions were significantly different (adjusted *p* < 0.05; **Figure 7B**). Finally, Rrat-GPP-9426 one-way ANOVA showed a significant effect (F(4, 10) = 199.5, *p* < 0.05). Homogeneity of variance was met (Brown–Forsythe: F(4, 10) = 0.695, *p* = 0.621). Dunnett’s post-hoc test, comparing each dilution against the negative control, indicated that all dilutions were significantly different (adjusted *p* < 0.05; **Figure 7C**). Semi-quantification capabilities on gDNA for *R. exulans*, *R. norvegicus* and *R. rattus* were R^2^ = 0.88, 0.96 and 0.94, respectively (**Figure 7**).

**Figure 7.**
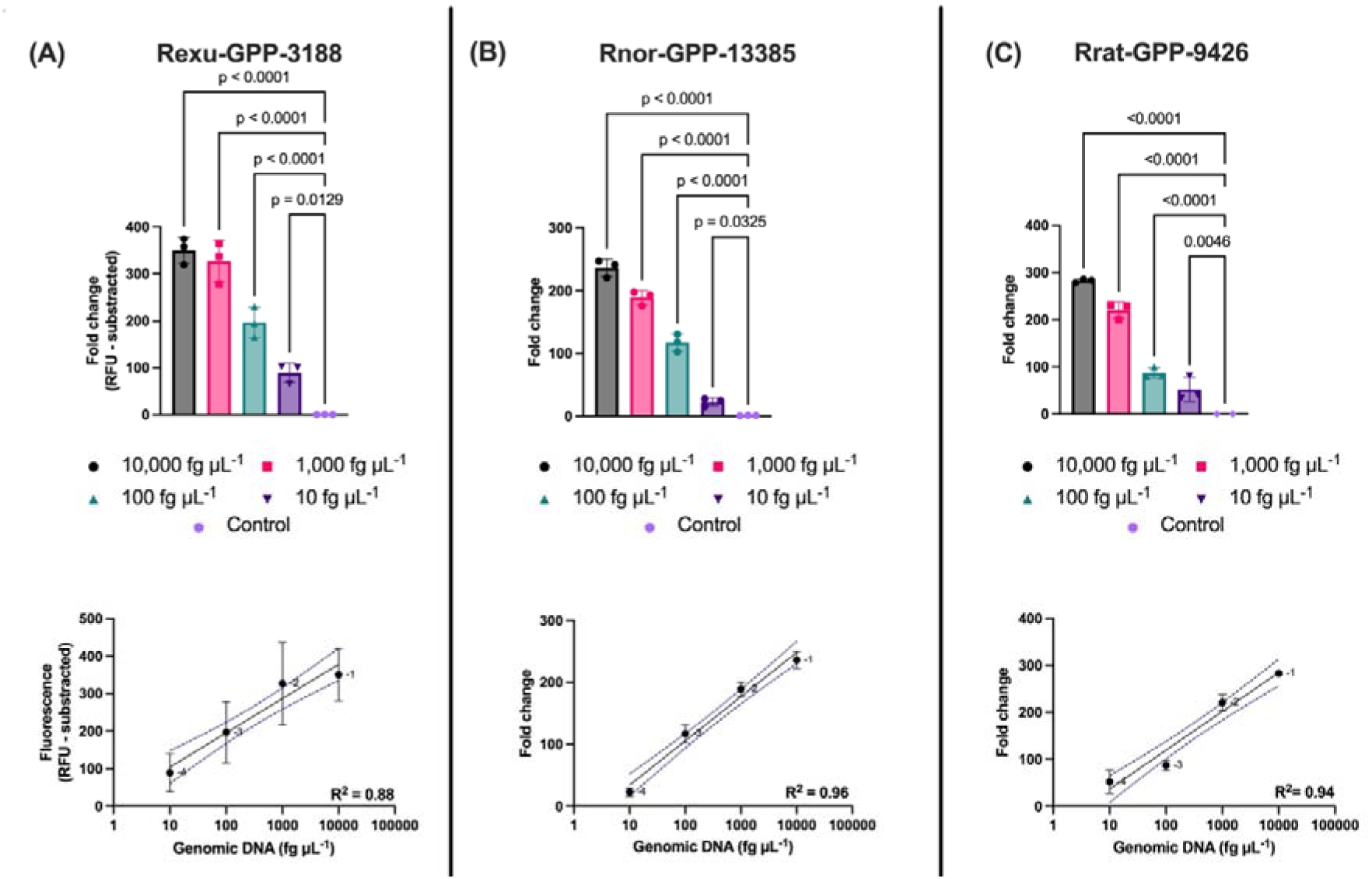
SENTINEL guide-primer pair semi-quantitative window in genomic DNA. **(A)** Rexu-GPP-3188 fold change on gDNA one-way ANOVA was significant, and Dunnett’s post-hoc test, comparing each dilution against the negative control, indicated that all GPPs were significantly different (adjusted p < 0.05). **(B)** Rnor-GPP13385 fold change on gDNA one-way ANOVA was significant and Dunnett’s post-hoc test, comparing each dilution against the negative control, indicated that all GPPs were significantly different (adjusted p < 0.05). **(C)** Rrat-GPP-9426 fold change on gDNA one-way ANOVA was significant and Dunnett’s post-hoc test, comparing each dilution against the negative control, indicated that all GPPs were significantly different (adjusted p < 0.05). All dilutions were labelled accordingly in the interpolation graphs (-1, -2, -3 and -4). Abbreviations: GPP: guide-primer pair; Rexu: *Rattus exulans*; Rnor: *Rattus norvegicus*; Rrat: *Rattus rattus*; SENTINEL: Smart Environmental Nucleic-acid Tracking using Inference from Neural-networks for Early-warning Localization.

### SENTINEL GPPs validation for Rattus exulans and Rattus rattus using environmental DNA

As a final step, we intended to challenge and validate our SENTINEL platform GPPs with real eDNA samples. SENTINEL *R. exulans* assay using Rexu-GPP-3188 assay was successful (**Figure 8A**), detecting at least one positive on 8/8 *R.exulans* eDNA qPCR positives samples, without any cross-reactivity observed in 3/3 true eDNA qPCR negatives samples and the NTC (**Figure 8B**). The Rexu-GPP-3188 A_t_ on eDNA samples ranged from 20.51 to 54.30 min, with a mean A_t_ equal to 34.45 min **(Figure 8C).** Sample Cq values from qPCR benchmarking are found in **Supplemental Table 6.**

**Figure 8.**
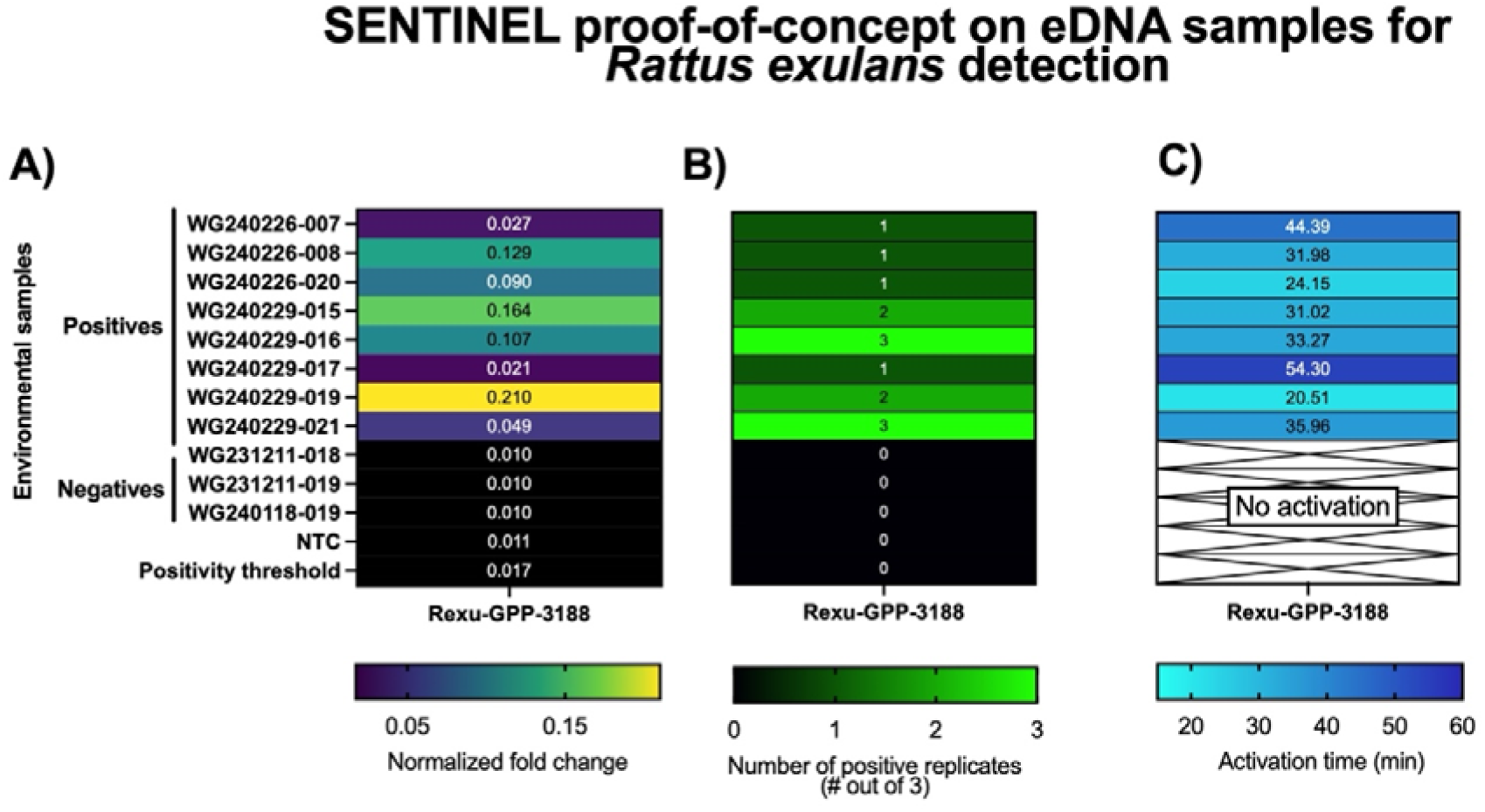
SENTINEL Rexu-GPP-3188 validation on environmental samples. **(A)** Normalized fold change is shown for all samples. Positivity threshold is also shown for reference. **(B)** SENTINEL positive calls above the threshold. (C) Activation time of the positive samples shown in **A** and **B**. All boxes represent the mean when possible. Abbreviations: eDNA: environmental DNA; Rexu: *Rattus exulans*; GPP: guide-primer pair; NTC: non-template control; SENTINEL: Smart Environmental Nucleic-acid Tracking using Inference from Neural-networks for Early-warning Localization.

Similar results were achieved with SENTINEL *R. rattus* assay using Rrat-GPP-9426 (**Figure 9A**), detecting at least one positive on 7/7 *R. rattus* eDNA samples that have positives hits in qPCR and metabarcoding (**Supplemental Table 6**). No cross-reactivity was observed in the NTC and true eDNA negatives also benchmarked with qPCR and metabarcoding (**Figure 9B**). The Rrat-GPP-9426 A_t_ on eDNA samples ranged from 15.44 to 43.34 min, with a mean A_t_ equal to 28.65 min (**Figure 9C**).

**Figure 9.**
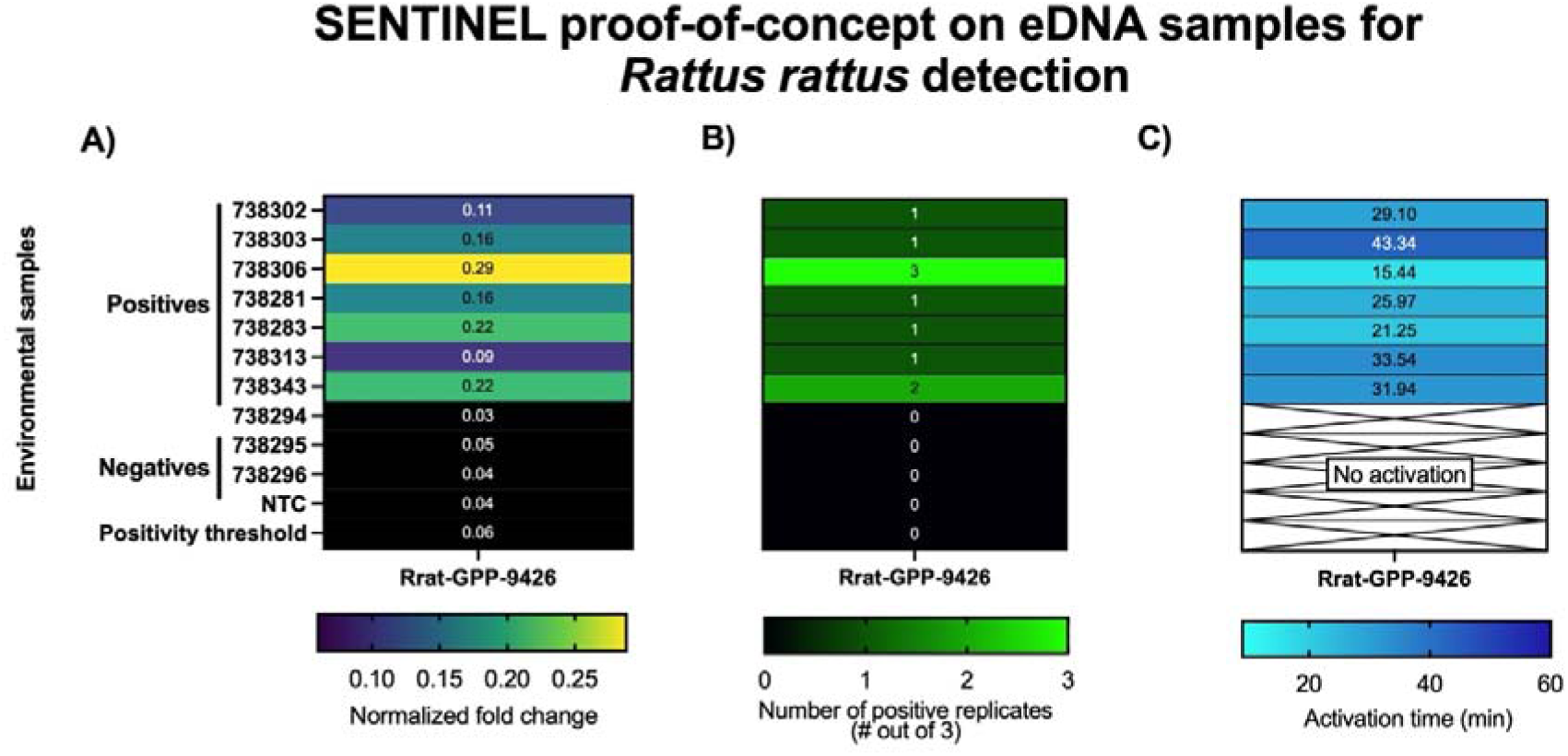
SENTINEL Rrat-GPP-9426 validation on environmental samples. **(A)** Normalized fold change is shown for all samples. Positivity threshold is also shown for reference. **(B)** SENTINEL positive calls above the threshold. **(C)** Activation time of the positive samples shown in **A** and **B**. All boxes represent the mean when possible. Abbreviations: eDNA: environmental DNA; Rrat: *Rattus rattus*; GPP: guide-primer pair; NTC: non-template control; SENTINEL: Smart Environmental Nucleic-acid Tracking using Inference from Neural-networks for Early-warning Localization.

## Discussion

Our study focused on deploying a SENTINEL proof of concept for *R. exulans*, *R. norvegicus*, and *R. rattus* using gDNA reference samples from the US and NZ. Demonstrating feasibility at the gDNA level provides a critical foundation for subsequent optimization and future deployment of field-based tools for eDNA detection, where environmental complexity and lower template availability might present additional challenges.

Our first step was to deploy a pre-trained artificial intelligence model to explore the full mitochondrial genomes of the target species for GPP identification (**Supplemental Table 1**). We obtained candidate GPPs for each target *Rattus* species. (**Table 1**). The best *in silico*-predicted candidates were selected and subsequently tested using 10 pM plasmid DNA and a final gDNA concentration of 0.01 ng µL^-1^. For further metric analysis to facilitate GPP selection, we have described A_t_, inspired by the calculations of Metsky et al. (2022). This metric may help provide an exact number to whether a GPP exhibits quick early activation above an arbitrary threshold. We believe A_t_ may be critical to provide extra information to understand when samples are already positive and a decision can already be called, especially in cases where some invasive species should not be present in a given site or sample.

Rexu-GPP-3188, Rnor-GPP-13385 and Rrat-GPP-9426 were selected as the final GPP candidates for detecting the corresponding species with high endpoint fluorescence and the lowest A_t_. Interestingly, each GPP performed considerable better on a different reference gDNA sample with exception of Rrat-GPP-9426 (**see Figure 2**, **Figure 3 and Figure 4**). This could indicate that either each GPP may have a higher preference to certain population or that less mitochondrial genomes were obtained during gDNA extraction and purification, directly impacting A_t_ and endpoint fluorescence. We believe that to understand the variation in detection rates with genetic diversity of these species, we suggest a robust study testing these assays across multiple populations is necessary to enable a detection of all *Rattus rattus* invasive populations, for example.

In this study, US and NZ gDNA samples were used as a representation of potential genomic variability across the same species but from different locations (see **Supplemental Table 4**). Nevertheless, on every GPP tested, the SENTINEL platform could detect both, US and NZ references, although some differences were clearly observed in endpoint fluorescence and A_t_ (**Figure 2; Figure 3**), which may be linked to mitochondrial DNA depletion, DNA degradation, variations in genetic diversity, or variations in DNA extraction methods.

After screening GPPs functionality over plasmid DNA and gDNA, we moved into testing all GPPs cross-reactivity with a panel of 11 different gDNAs of closely related rat species from reference samples from US and NZ. In short, no GPP had a fold change above the positive threshold set (**Supplemental Table 7**). Hence, these results indicate and support that the SENTINEL platform may be highly specific for detecting closely related rat species (**Figure 5**).

To further challenge our proof-of-concept validation on closely rat species and explore whether the SENTINEL platform is reproducible with previous results (Durán-Vinet et al. 2025), we tested the sensitivity of candidate GPPs in plasmid DNA and gDNA. As observed in **Figure 6**, GPPs exhibit a significant difference compared to the control in plasmid DNA, with an average R^2^ score of ∼0.73, which aligns with previous results of the SENTINEL platform, but in marine invasive species (Durán-Vinet et al. 2025). Similar results were obtained with gDNA (**Figure 7**). However, candidate GPPs increased their R^2^ values to an average of ∼0.93. These results are interesting as gDNA is a more complex sample than plasmid DNA, where we expected that gDNA complexity would decrease the efficiency of the reaction, however, semi-quantification efficiency was increased. Nevertheless, these results further underpin the SENTINEL’s platform semi-quantitative capabilities that could be of use to potentially broadly understand target abundance.

Interestingly, when these results are put into context, a single rat cell has ∼2.75Gbps (Howe et al. 2021), which in a -4 dilution (equal to 0.00001 ng uL^-1^) is similar to ∼5.5 aM or ∼5 copies µL_1_ mitochondrial DNA-equivalent cells mL^-1^ ∼0.0017-0.007 cells mL^-1^ (assuming a cell might have between 500∼2000 mitochondria per cell). Therefore, we can infer that SENTINEL can be highly specific within closely related species, but also an ultra-sensitive platform for CRISPR-eBx deployment contexts.

Further validation was then undertaken on obtained eDNA samples from US (*R. exulans*) and NZ (*R. rattus*) as a final proof of concept. We observed 100% agreement between qPCR and metabarcoding (**Supplemental Table 6**) with the positive calls with SENTINEL (**Figure 8A; Figure 9A**). It was also observed that SENTINEL had stochasticity across all tested samples (**Figure 8B; Figure 9B)**, which could explain due to the low abundance and fragmented nature of eDNA (Power et al. 2023; West and Deagle 2025). Further, large-scale standardized validation is required to fully understand this effect weight.

In the same context of eDNA low abundance and fragmentation, and potential high diverse of other taxa present in an eDNA sample (Durán-Vinet et al. 2025b), the A_t_ of eDNA samples was much higher than plasmid and gDNA (**Figure 8C; Figure 9C**). This means that the overall efficiency of the reaction is under stress, therefore, slower. Moreover, looking at plasmid DNA and gDNA results (e.g., **Figure 3**) and eDNA results (e.g., **Figure 8**), there is a clear pattern that sample complexity increases from plasmid to eDNA, as the endpoint fluorescence decreases and A_t_ increases substantially. Furthermore, eDNA samples only reached ∼20% of the observed gDNA performance at best. This is a critical point for CRISPR-eBx assay validations, where high gDNA performance must be sought prior eDNA deployment, as there might be a substantial performance drop that is unlikely to be linked to ‘primer’ performance, but rather the high complexity of eDNA samples.

## Concluding remarks

In this study, we successfully deployed the SENTINEL platform to detect invasive rat species using gDNA from three closely related, invasive rat species: *R. exulans*, *R. norvegicus* and *R. rattus*. By integrating a previous workflow based on a AI-assisted GPP design with a proven systematic experimental validation, we identified and refined a subset of high-performing GPPs capable of discriminating between closely related rodent taxa. These assays demonstrated strong and consistent endpoint fluorescence, robust activation time for *R. exulans*, *R. norvegicus* and *R. rattus*.

Overall, the current obtained results highlights SENTINEL’s capacity to deliver highly sensitive and highly specific detection of closely related invasive rats using gDNA and eDNA. This represents an important step forward for CRISPR-eBx for land biosecurity applications, where rapid and species-resolved detection is critical for early intervention, population management and decrease their wide economic impact.

In summary, this study further demonstrates the feasibility of SENTINEL as a robust CRISPR-eBx application for detecting invasive land species that could be closely related. The pilot deployment on eDNA for *R. exulans* and *R. rattus* provides a strong empirical foundation and further confirms the utility and reproducibility of the SENTINEL platform as an AI-guided development and pre-selection of targets for complex biosecurity scenarios, such as closely related land invasive species.

## Author contributions

**Benjamín Durán-Vinet**: Conceptualization, Methodology, Investigation, Formal analysis, Data curation, Visualization, Project administration, Writing – original draft, Writing – review & editing. **Antoinette J. Piaggio**: Conceptualization, Methodology, Resources, Funding acquisition, Writing – review & editing. **Nicholas Foster**: Resources, Validation. **Anna Clark**: Resources, Writing – review & editing. **Madison Sayler**: Investigation, Data, Resources. **Gert-Jan Jeunen**: Methodology, Supervision, Writing – review & editing. **Jackson Treece**: Methodology, Investigation, Resources, Validation. **Sara Ferreira**: Methodology, Investigation, Resources. **Catherine Collins**: Resources, Investigation, Writing – review & editing. **Stacey Buckelew**: Investigation, Resources, Project administration, Funding acquisition, Writing – review & editing. **Neil J. Gemmell**: Conceptualization, Supervision, Project administration, Funding acquisition, Writing – review & editing.

## Acknowledgements

Ministry of Business, Innovation, and Employment (MBIE) project: A toolbox to underpin and enable tomorrow’s marine biosecurity system (MBIE CAWX1904) funded the cost for this study and B. D.-V. PhD scholarship. Financial support was provided by the US Air Force Pacific Air Forces and the 611th Civil Engineer Squadron through the Wake Atoll Rat Eradication Project. The funders had no role in study design, data collection nor analysis. This research was also supported in part by the U.S. Department of Agriculture, Animal Plant Health Inspection Service (APHIS), Wildlife Services, National Wildlife Research Center (NWRC) with funding through the US Fish and Wildlife Invasive Species program. The findings and conclusions in this publication are those of the author(s) and should not be construed to represent the views of the USDA or U.S. government.

