## Supplemental for "A streamlined proof of concept CRISPR-based environmental biosurveillance platform for the detection of closely related invasive rats"

*Original article; supplemental information*

**Supplemental Methods**

To pursue a more metric-based decision making for GPP selection beyond endpoint fluorescence, our team has described an additional useful metrics mathematically calculated that can be easily computed from fluorescence kinetics on positive controls, either plasmid DNA, genomic DNA (gDNA) and also environmental DNA (eDNA) as shown in the results of this article.

Firstly, we setup the baseline values, with SENTINEL platform assays data from no-template controls (NTCs) as sample to obtain SENTINEL fluorescence noise for each GPP. Accordingly, to obtain a stable endpoint baseline mean ($\bar{B}_{endpoint}$), we used the last five timepoints (*t*) of the average subtracted fluorescence of the three replicates (**Equation 1**). Using the last five timepoints ensures that we are obtaining a representative sample of the end portion of the reaction. This is especially important when last time points might or might not have reached plateau. The average of these values was also used to obtain their standard deviation (STD), and then sum it up to $\bar{B}_{endpoint}$ to use it as $B_{reference}$ (**Equation 2**) for fold-change calculation for their corresponding GPP.

$$\bar{B}_{endpoint}= \frac{1}{5}\sum_{t=60}^{60} B_{t}$$

**Equation 1. Mean of the baseline endpoint subtracted fluorescence.**

The *t* value is equal to the subtracted fluorescence in *t* timepoint. Units are in RFU.

$$\bar{B}_{reference}= \bar{B}_{endpoint}+\bar{B}_{STD}$$

**Equation 2. Baseline reference for fold-change calculation.**

STD: Standard deviation of the sample.

Then, $\bar{B}_{reference}$ is multiplied by an arbitrary threshold multiplier (${Ar}_{tm}$) that depends on each GPP overall performance (we used in this study values from 3 to 8). Higher thresholds make calculations more conservative, while lower may increase assay sensitivity but also increase potential false positives. This value is indexed as fluorescence threshold ($F_{thr}$) that is used as a reference point to indicate when the subtracted fluorescence of a single replicate ($F_{t}$) or average ($\bar{F}_{t}$) in a given *t* is above $F_{thr}$ known as activation time ($A_{t}$).

$A_{t}$ is calculated using *t* < $F_{thr}$ ($t_{before}$) and *t* > $F_{thr}$ (t_after_); and $F_{t}$ < $F_{thr}$ ($F_{t,before}$) and $F_{t}$ > $F_{thr}$ ($F_{t, after}$) (**Equation 3**).

$$A_{t}= \frac{{[t}_{before}-\left( t_{after}-t_{before} \right)]*{(F}_{thr}-F_{t,before})}{F_{t,after}-F_{thr}}$$

**Equation 3. Activation time calculation.**

Units are in min.

Fold change is calculated using raw fluorescence or mean raw fluorescence readings (usually endpoint) and subtracting and dividing using $\bar{B}_{endpoint}$ (**equation 4**). This ensures that the background noise is subtracted from the sample.

$$Fold change=\frac{(F- \bar{B}_{endpoint})}{\bar{B}_{endpoint}}$$

**Equation 4. Fold change calculation.**

No units.

**Supplemental tables**

**Supplemental Table 1. Reference accession numbers.**

| **Species name** | **GenBank accession number (on target)** | **GenBank accession number (off target)** |
| --- | --- | --- |
| *Rattus exulans* | EU273709; EU273710; EU273711; KJ530564; KY814709; KY814710; KY814711; KY814712; KY814713; KY814714; KY814715; KY814716; KY814717; KY814718; KY814719; KY814720; KY814721; MN126569; NC_012389; OM908890; OM908891; OM908892; OM908893; OM908894; OM908895; OM908896; OM908897; OM908898; OM908899; OM908900; OM908901; OM908902 | AY172581; AY584828; AY769440; CM080009; DQ124371; DQ124372; DQ124373; DQ124374; DQ124375; DQ124376; DQ673907; DQ673908; DQ673909; DQ673910; DQ673911; DQ673912; DQ673913; DQ673914; DQ673915; DQ673916; DQ673917; EU273708; EU450583; FJ355927; FJ919759; FJ919760; FJ919761; FJ919762; FJ919763; FJ919764; FJ919766; FJ919767; FJ919768; FJ919769; FJ919770; FJ919771; GU570659; GU570660; GU570661; GU570662; GU570663; GU570664; GU570665; GU997608; GU997609; GU997610; GU997611; HM152027; HM152028; HQ439465; HQ439466; HQ439469; HQ439470; HQ439486; HQ439492; HQ586004; JN398399; JN398400; JQ003190; JX105355; JX105356; KC152486; KF011916; KF011917; KF937876; KJ530565; KJ939360; KJ939361; KM009112; KM009113; KM009114; KM114603; KM114604; KM114605; KM114606; KM114607; KM114608; KM577634; KM577635; KM657952; KM657953; KM820831; KM820832; KM820833; KM820834; KM820835; KM820837; KP099715; KP100657; KP233827; KP233832; KP241960; KP244683; KP876560; KU200226; KU745736; KX058347; KY117577; KY117578; KY117579; KY117580; KY117581; KY464180; KY611359; KY611360; KY611361; KY611362; KY611363; KY611364; KY611365; KY611366; KY611367; KY611368; KY611369; KY611370; KY611371; KY611372; KY611373; KY611374; KY611375; KY611376; KY611377; KY611378; KY611379; KY611380; KY611381; KY611382; KY611383; KY611384; KY611385; KY611386; KY611387; KY611388; KY611389; KY611390; KY707300; MG182016; MN126561; MN126562; MN126563; MN126564; MN126566; MN126567; MN126568; MN964117; MT259589; MT259590; MT410886; MT862372; MT937073; MUSMTDNAD; MW209724; MW209726; MZ014548; NC_001665; NC_005089; NC_011638; NC_012374; NC_012461; NC_014855; NC_014858; NC_014861; NC_014864; NC_014867; NC_014871; NC_023347; NC_029888; NC_033356; NC_035594; NC_039670; NC_040919; NC_046686; NC_049040; NC_049042; NC_068809; NC_068811; OK054583; OM574930; OM574931; OM574932; OM574933; OM574934; OM574935; OM574936; OM574937; OM574938; OM574939; OM574940; OM574941; OM574942; OM574943; OM574944; OM574945; OM574946; OM574947; OM574948; OM574949; OM574950; OM574951; OM574952; OM574953; OM574954; OM574955; OM574956; OM574957; OM574958; OM574959; OM574960; OM574961; OM574962; OM574963; OM574964; OM574965; OM574966; OM574967; OM574968; OM574969; OM574970; OM948981; OM963003; ON528109; OP251025; OP859007; OP859008; OP859009; OP859010; OP859011; OP859012; OR077806; OR077807; OR077808; OR077854; OR077855; OR077856; OR077857; OR085738; OR840749; OW971719; OW971866; OX389814; OX439034; PP454668; PP454669; PP454688; PP454689; PP454690; PP454691; PP454692 |
| *Rattus norvegicus* | AY172581; AY769440; DQ673907; DQ673908; DQ673909; DQ673910; DQ673911; DQ673912; DQ673913; DQ673914; DQ673915; DQ673916; DQ673917; FJ919759; FJ919760; FJ919761; FJ919762; FJ919763; FJ919764; FJ919766; FJ919767; FJ919768; FJ919769; FJ919770; FJ919771; GU997608; GU997609; GU997610; GU997611; HM152027; HM152028; JX105355; JX105356; KF011917; KJ530565; KJ939360; KJ939361; KM009112; KM009113; KM009114; KM114603; KM114604; KM114605; KM114606; KM114607; KM114608; KM577634; KM577635; KM657952; KM657953; KM820831; KM820832; KM820833; KM820834; KM820835; KM820837; KP099715; KP100657; KP233827; KP233832; KP241960; KP244683; MW209726; MZ014548; NC_001665 | AY584828; CM080009; DQ124371; DQ124372; DQ124373; DQ124374; DQ124375; DQ124376; EU273708; EU273709; EU273710; EU273711; EU450583; FJ355927; GU570659; GU570660; GU570661; GU570662; GU570663; GU570664; GU570665; HQ439465; HQ439466; HQ439469; HQ439470; HQ439486; HQ439492; HQ586004; JN398399; JN398400; JQ003190; KC152486; KF011916; KF937876; KJ530564; KP876560; KU200226; KU745736; KX058347; KY117577; KY117578; KY117579; KY117580; KY117581; KY464180; KY611359; KY611360; KY611361; KY611362; KY611363; KY611364; KY611365; KY611366; KY611367; KY611368; KY611369; KY611370; KY611371; KY611372; KY611373; KY611374; KY611375; KY611376; KY611377; KY611378; KY611379; KY611380; KY611381; KY611382; KY611383; KY611384; KY611385; KY611386; KY611387; KY611388; KY611389; KY611390; KY707300; KY814709; KY814710; KY814711; KY814712; KY814713; KY814714; KY814715; KY814716; KY814717; KY814718; KY814719; KY814720; KY814721; MG182016; MN126561; MN126562; MN126563; MN126564; MN126566; MN126567; MN126568; MN126569; MN964117; MT259589; MT259590; MT410886; MT862372; MT937073; MUSMTDNAD; MW209724; NC_005089; NC_011638; NC_012374; NC_012389; NC_012461; NC_014855; NC_014858; NC_014861; NC_014864; NC_014867; NC_014871; NC_023347; NC_029888; NC_033356; NC_035594; NC_039670; NC_040919; NC_046686; NC_049040; NC_049042; NC_068809; NC_068811; OK054583; OM574930; OM574931; OM574932; OM574933; OM574934; OM574935; OM574936; OM574937; OM574938; OM574939; OM574940; OM574941; OM574942; OM574943; OM574944; OM574945; OM574946; OM574947; OM574948; OM574949; OM574950; OM574951; OM574952; OM574953; OM574954; OM574955; OM574956; OM574957; OM574958; OM574959; OM574960; OM574961; OM574962; OM574963; OM574964; OM574965; OM574966; OM574967; OM574968; OM574969; OM574970; OM908890; OM908891; OM908892; OM908893; OM908894; OM908895; OM908896; OM908897; OM908898; OM908899; OM908900; OM908901; OM908902; OM948981; OM963003; ON528109; OP251025; OP859007; OP859008; OP859009; OP859010; OP859011; OP859012; OR077806; OR077807; OR077808; OR077854; OR077855; OR077856; OR077857; OR085738; OR840749; OW971719; OW971866; OX389814; OX439034; PP454668; PP454669; PP454688; PP454689; PP454690; PP454691; PP454692 |
| *Rattus rattus* | FJ355927; MT862372; MW209724; NC_012374; OM574930; OM574931; OM574932; OM574933; OM574934; OM574935; OM574936; OM574937; OM574938; OM574939; OM574940; OM574941; OM574942; OM574943; OM574944; OM574945; OM574946; OM574947; OM574948; OM574949; OM574950; OM574951; OM574952; OM574953; OM574954; OM574955; OM574956; OM574957; OM574958; OM574959; OM574960; OM574961; OM574962; OM574963; OM574964; OM574965; OM574966; OM574967; OM574968; OM574969; OM574970 | AY172581; AY584828; AY769440; CM080009; DQ124371; DQ124372; DQ124373; DQ124374; DQ124375; DQ124376; DQ673907; DQ673908; DQ673909; DQ673910; DQ673911; DQ673912; DQ673913; DQ673914; DQ673915; DQ673916; DQ673917; EU273708; EU273709; EU273710; EU273711; EU450583; FJ919759; FJ919760; FJ919761; FJ919762; FJ919763; FJ919764; FJ919766; FJ919767; FJ919768; FJ919769; FJ919770; FJ919771; GU570659; GU570660; GU570661; GU570662; GU570663; GU570664; GU570665; GU997608; GU997609; GU997610; GU997611; HM152027; HM152028; HQ439465; HQ439466; HQ439469; HQ439470; HQ439486; HQ439492; HQ586004; JN398399; JN398400; JQ003190; JX105355; JX105356; KC152486; KF011916; KF011917; KF937876; KJ530564; KJ530565; KJ939360; KJ939361; KM009112; KM009113; KM009114; KM114603; KM114604; KM114605; KM114606; KM114607; KM114608; KM577634; KM577635; KM657952; KM657953; KM820831; KM820832; KM820833; KM820834; KM820835; KM820837; KP099715; KP100657; KP233827; KP233832; KP241960; KP244683; KP876560; KU200226; KU745736; KX058347; KY464180; KY611359; KY611360; KY611361; KY611362; KY611363; KY611364; KY611365; KY611366; KY611367; KY611368; KY611369; KY611370; KY611371; KY611372; KY611373; KY611374; KY611375; KY611376; KY611377; KY611378; KY611379; KY611380; KY611381; KY611382; KY611383; KY611384; KY611385; KY611386; KY611387; KY611388; KY611389; KY611390; KY707300; KY814709; KY814710; KY814711; KY814712; KY814713; KY814714; KY814715; KY814716; KY814717; KY814718; KY814719; KY814720; KY814721; MG182016; MN126561; MN126562; MN126563; MN126564; MN126566; MN126567; MN126568; MN126569; MN964117; MT259589; MT259590; MT410886; MT937073; MUSMTDNAD; MW209726; MZ014548; NC_001665; NC_005089; NC_011638; NC_012389; NC_012461; NC_014855; NC_014858; NC_014861; NC_014864; NC_014867; NC_014871; NC_023347; NC_029888; NC_033356; NC_035594; NC_039670; NC_040919; NC_046686; NC_049040; NC_049042; NC_068809; NC_068811; OK054583; OM908890; OM908891; OM908892; OM908893; OM908894; OM908895; OM908896; OM908897; OM908898; OM908899; OM908900; OM908901; OM908902; OM948981; OM963003; ON528109; OP251025; OP859007; OP859008; OP859009; OP859010; OP859011; OP859012; OR077806; OR077807; OR077808; OR077854; OR077855; OR077856; OR077857; OR085738; OR840749; OW971719; OW971866; OX389814; OX439034; PP454668; PP454669; PP454688; PP454689; PP454690; PP454691; PP454692 |

**Supplemental Table 2. Command lines used in ADAPT for each pipeline deployment.**

| Target species | Command line |
| --- | --- |
| *Rattus exulans* | design.py complete-targets fasta ./Designs/Input/Rexulans_ontarget.fasta -o ./Designs/Output/R.exulans_designsv2 --obj maximize-activity --specific-against-fastas ./Designs/Input/Off-target_vs_R.exulans.fasta --id-m 4 --id-frac 0.01 -gl 28 -gm 0 -pl 30 -pm 3 -pp 0.98 --primer-gc-content-bounds 0.30 0.70 --max-target-length 300 --maximization-algorithm random-greedy --predict-cas13a-activity-model --best-n-targets 10 --seed 104 --verbose |
| *Rattus norvegicus* | design.py complete-targets fasta ./Designs/Input/Rnorvegicus_ontarget.fasta -o ./Designs/Output/R.norvegicus_designsv1 --obj maximize-activity --specific-against-fastas ./Designs/Input/Off-target_vs_R.norvegicus.fasta --id-m 4 --id-frac 0.01 -gl 28 -gm 0 -pl 30 -pm 3 -pp 0.98 --primer-gc-content-bounds 0.30 0.70 --max-target-length 300 --maximization-algorithm random-greedy --predict-cas13a-activity-model --best-n-targets 50 --seed 105 --verbose |
| *Rattus rattus* | design.py complete-targets fasta ./Designs/Input/Rrattus_ontarget.fasta -o ./Designs/Output/R.rattus_designsv3 --obj maximize-activity --specific-against-fastas ./Designs/Input/Off-target_vs_R.rattus.fasta --id-m 4 --id-frac 0.01 -gl 28 -gm 0 -pl 30 -pm 3 -pp 0.98 --primer-gc-content-bounds 0.30 0.70 --max-target-length 300 --maximization-algorithm random-greedy --predict-cas13a-activity-model --best-n-targets 10 --seed 106 --verbose |

**Supplemental Table 3. Consensus sequence for plasmid insert.**

| Target species | Guide-primer pair | Insert sequence |
| --- | --- | --- |
| *Rattus exulans* | Rexu-GPP-3188 | ATTCAAAATACTCTCTATTTGGAGCTCTACGAGCCGTCGCCCAAACTATCTCTTACGAAGTTACAATAGCCATCATCCTACTATCTGTACTTTTAATAAATGGCTCTTTCTCCCTACAAATACTTATCACCACACAAGAACACATCTGAC |
| *Rattus norvegicus* | Rnor-GPP-1555 | ACCATCAGAACAACAAATCAAAATGTAAACTTAAAATATAGCCAAAAGAGGGACAGCTCTTTAGGAAAAGGAAAAAACCTTAAATAGTGAATAAACAACTACAATCACTTAACCATTGTAGGCTTAAAAGCAGCCATCAATAAAGAAAGC |
|  | Rnor-GPP-13385 | TCAATAAGCAAACCCACCAAACTATCATCATTCTCAACCTCCCTAGGCTACTACCCACCAATTATACACCGAATTATTCCTCAAAAAACTCTAAATTCTAGCTACAAATTATCCTTAAACCTACTAGACCTAATCTGACTAGAAAAGACA |
| *Rattus rattus* | Rrat-GPP-4487 | GGAGGATTAAACCAAACACAAACACGGAAGATCATAGCATACTCATCAATTGCTCATATAGGATGAATAGTGGCAATTCTCCCCCACAATCCTAATCTGACACTCCTAAACCTAACAATTTATATCCTACTCACTATCCCAATATTCACC |
|  | Rrat-GPP-9426 | AACACCATTTTGGGTTTGAAGCCGCAGCATGATATTGACATTTCGTAGATGTAGTTTGACTATTCCTATATGTTTCTATCTATTGATGAGGATCATACTCCCTTAGTATAATCAATACAACTGACTTCCAATCAGTAAATTCTGAAAAACCCAGAAGAGAGTAATTAACCTGTTTATTATTATTACAATCAATACCACCCTCTCTCTCATTCTTGCTTCAATCGCATTCT |

Abbreviations: GPP: guide-primer pair; Rexu: *Rattus exulans*; Rnor: *Rattus norvegicus*; Rrat: *Rattus rattus*.

**Supplemental Table 4**. All species used as reference samples in this study.

| Species name | Origin | Provided as | Provided by | Institution |
| --- | --- | --- | --- | --- |
| *Homo sapiens* | Human cell line | Purified gDNA | Dr Indranil Basak | University of Otago, Department of Biochemistry New Zealand |
| *Mus musculus* | United States | Purified gDNA | Dr Antoinette J. Piaggio | USDA, United States |
| *Neotoma albigula* | United States | Purified gDNA | Dr Antoinette J. Piaggio | USDA, United States |
| *Rattus exulans* | United States | Purified gDNA | Dr Antoinette J. Piaggio | USDA, United States |
| *Rattus norvegicus* | United States | Purified gDNA | Dr Antoinette J. Piaggio | USDA, United States |
| *Rattus rattus* | United States | Purified gDNA | Dr Antoinette J. Piaggio | USDA, United States |
| *Rattus tanezumi* | United States | Purified gDNA | Dr Antoinette J. Piaggio | USDA, United States |
| *Mus musculus* | New Zealand | Purified gDNA | Dr Victoria J Sugrue | University of Otago, Department of Anatomy, New Zealand |
| *Rattus exulans* | New Zealand | Purified gDNA | Dr Catherine Collins | University of Otago, Department of Anatomy, New Zealand |
| *Rattus norvegicus* | New Zealand | Purified gDNA | Dr Catherine Collins | University of Otago, Department of Anatomy, New Zealand |
| *Rattus rattus* | New Zealand | Purified gDNA | Dr Anna Clark | University of Otago, Department of Anatomy, New Zealand |

**Supplemental Table 5. Tested dilutions for SENTINEL sensitivity and cross-reactivity testing**

| Dilution ID | Input^a^  (fg µL^-1^) | Final concentration^b^  (fg µL^-1^) | Total input  (fg reaction^-1^) | Use |
| --- | --- | --- | --- | --- |
| -1 | ~100,000 | ~10,000 | ~200,000 | Sensitivity  Cross-reactivity |
| -2 | ~10,000 | ~1,000 | ~20,000 | Sensitivity |
| -3 | ~1,000 | ~100 | ~2,000 | Sensitivity |
| -4 | ~100 | ~10 | ~200 | Sensitivity |
| -5 | ~10 | ~1 | ~20 | Sensitivity |

^a^ Sample input is always 2 µL at the indicated concentration.

^b^ Final reaction concentration. Input concentration is diluted a 1:10 ratio to provide the final concentration.

Abbreviations: SENTINEL: Smart Environmental Nucleic-acid Tracking using Inference from Neural-networks for Early-warning Localization.

**Supplemental Table 6. Environmental DNA samples information**

| Sample ID | qPCR /metabarcoding identification call | Use | Location (place, country) | N of positive replicates | Cq mean or quantification mean (cp/µL) | Metabarcoding reads | Reference |
| --- | --- | --- | --- | --- | --- | --- | --- |
| WG240226-007 | *R. exulans* (qPCR) | True eDNA positive validation | Wake Island (US) | 1/3 | 2.40 cp/µL | NA | Toni? |
| WG240226-008 | *R. exulans* (qPCR) | True eDNA positive validation | Wake Island (US) | 3/3 | 4.18 cp/µL | NA |  |
| WG240226-020 | *R. exulans* (qPCR) | True eDNA positive validation | Wake Island (US) | 3/3 | 2.65 cp/µL | NA |  |
| WG240229-015 | *R. exulans* (qPCR) | True eDNA positive validation | Wake Island (US) | 3/3 | 7.74 cp/µL | NA |  |
| WG240229-016 | *R. exulans* (qPCR) | True eDNA positive validation | Wake Island (US) | 3/3 | 5.29 cp/µL | NA |  |
| WG240229-017 | *R. exulans* (qPCR) | True eDNA positive validation | Wake Island (US) | 3/3 | 3.68 cp/µL | NA |  |
| WG240229-018 | *R. exulans* (qPCR) | True eDNA positive validation | Wake Island (US) | 2/3 | 1.95 cp/µL | NA |  |
| WG240229-021 | *R. exulans* (qPCR) | True eDNA positive validation | Wake Island (US) | 3/3 | 3.41 cp/µL | NA |  |
| 783302 | *R. rattus (qPCR and metabarcoding)* | True eDNA positive validation | Barlow Creek - Manifold B (NZ) | 2/2 | 34.70 | 17866 | This study |
| 738303 | *R. rattus (qPCR and metabarcoding)* | True eDNA positive validation | Barlow Creek - Manifold B (NZ) | 2/2 | 34.45 | 10482 | This study |
| 738306 | *R. rattus (qPCR and metabarcoding)* | True eDNA positive validation | Barlow Creek - Manifold B (NZ) | 2/2 | 34.21 | 10680 | This study |
| 738281 | *R. rattus (qPCR and metabarcoding)* | True eDNA positive validation | Dale Creek - Manifold B (NZ) | 2/2 | 38.09 | 2106 | This study |
| 738283 | *R. rattus (qPCR and metabarcoding)* | True eDNA positive validation | Dale Creek - Manifold B (NZ) | 2/2 | 37.45 | 1401 | This study |
| 738313 | *R. rattus (qPCR and metabarcoding)* | True eDNA positive validation | Barlow Creek - Manifold A (NZ) | 2/2 | 36.07 | 2495 | This study |
| 738343 | *R. rattus (qPCR and metabarcoding)* | True eDNA positive validation | Dale Creek - Manifold A (NZ) | 2/2 | 38.66 | 2119 | This study |
| 738294 | *R. rattus (qPCR and metabarcoding)* | True eDNA negative validation | Kiwi Jack Creek - Manifold A (NZ) | 0/2 | 0.00 | 0.00 | This study |
| 738295 | *R. rattus (qPCR and metabarcoding)* | True eDNA negative validation | Kiwi Jack Creek - Manifold A (NZ) | 0/2 | 0.00 | 0.00 | This study |
| 738296 | *R. rattus (qPCR and metabarcoding)* | True eDNA negative validation | Kiwi Jack Creek - Manifold A (NZ) | 0/2 | 0.00 | 0.00 | This study |
| WG231211-018 | *Mus musculus* | True eDNA negative validation | Midway Island (US) | 1/3 | 0.35 cp/µL | NA | This study |
| WG231211-019 | *Mus musculus* | True eDNA negative validation | Midway Island (US) | 2/3 | 1.06 cp/µL | NA | This study |
| WG240118-019 | *Mus musculus* | True eDNA negative validation | Midway Island (US) | 2/3 | 0.49 cp/µL | NA | This study |

**Supplemental Table 7**. Metrics of candidate guide-primer pairs for SENTINEL

| ID | Target | B̄_endpoint_ | Ar_tm_ | F_thr_ | Fold change | A_t_ (min) | ${\bar{\mathbf{F}}}_{\mathbf{endpoint}}$  (RFU) |
| --- | --- | --- | --- | --- | --- | --- | --- |
| Rexu-GPP-3188 | Plasmid | 0.400 | 4 | 1.600 | 3 | 4.643 | 133.392 |
|  | gDNA-US |  |  |  |  | 16.397 | 119.609 |
|  | gDNA-NZ |  |  |  |  | 19.536 | 52.983 |
| Rnor-GPP-1555 | Plasmid | 0.624 | 3 | 1.872 | 2 | 3.200 | 129.136 |
|  | gDNA-US |  |  |  |  | 16.548 | 30.023 |
| Rnor-GPP-13385 | Plasmid |  |  |  |  | 1.654 | 136.882 |
|  | gDNA-US |  |  |  |  | 12.512 | 112.631 |
|  | gDNA-NZ |  |  |  |  | 7.752 | 157.427 |
| Rrat-GPP-4487 | Plasmid | 0.189 | 8 | 1.512 | 7 | 4.156 | 134.889 |
|  | gDNA-US |  |  |  |  | 13.905 | 102.539 |
| Rrat-GPP-9426 | Plasmid |  |  |  |  | 2.486 | 152.232 |
|  | gDNA-US |  |  |  |  | 7.144 | 143.881 |
|  | gDNA-NZ |  |  |  |  | 7.917 | 147.211 |

B̄_endpoint_, F_thr_, A_t_, $\bar{F}_{\mathrm{endpoint}}$ are provided as the triplicate mean, with three significant figures and rounded.

$\bar{F}_{\mathrm{endpoint}}$ is the mean of the subtracted endpoint fluorescence of the last five timepoints across triplicates minus the baseline mean of the last five timepoints.

Abbreviations: gDNA: Genomic DNA; GPP: guide-primer pair; SENTINEL: Smart Environmental Nucleic-acid Tracking using Inference from Neural-networks for Early-warning Localization.
